# A tethering mechanism underlies Pin1-catalyzed proline *cis-trans* isomerization at a noncanonical site

**DOI:** 10.1101/2024.07.19.604348

**Authors:** Christopher C. Williams, Jonathan Chuck, Paola Munoz-Tello, Douglas J. Kojetin

## Abstract

The prolyl isomerase Pin1 catalyzes the *cis-trans* isomerization of proline peptide bonds, a noncovalent post-translational modification that influences cellular and molecular processes, including protein-protein interactions. Pin1 is a two-domain enzyme containing a WW domain that recognizes phosphorylated serine/threonine-proline (pS/pT-P) canonical motifs and an enzymatic PPIase domain that catalyzes proline *cis-trans* isomerization of pS/pT-P motifs. Here, we show that Pin1 uses a tethering mechanism to bind and catalyze proline *cis-trans* isomerization of a noncanonical motif in the disordered N-terminal activation function-1 (AF-1) domain of the human nuclear receptor PPARγ. NMR reveals multiple Pin1 binding regions within the PPARγ AF-1, including a canonical motif (pS112-P113) that when phosphorylated by the kinase ERK2 binds the Pin1 WW domain with high affinity. NMR methods reveal that Pin1 also binds and accelerates *cis-trans* isomerization of a noncanonical motif containing a tryptophan-proline motif (W39-P40) previously shown to be involved in an interdomain interaction with the C-terminal ligand-binding domain (LBD) of PPARγ. Cellular transcription studies combined with mutagenesis and Pin1 inhibitor treatment reveal a functional role for Pin1-mediated acceleration of *cis-trans* isomerization of the PPARγ W39-P40 motif. Our data inform a refined model of the Pin1 catalytic mechanism where the WW domain can bind a canonical pS/T-P motif and tether Pin1 to a target, which enables the PPIase domain to exert catalytic *cis-trans* isomerization at a distal noncanonical site.

**SIGNIFICANCE:** Pin1 is a multidomain prolyl isomerase enzyme that catalyzes the isomerization of proline peptide bonds, which naturally occur in *cis* and *trans* conformations that exchange on a timescale of seconds to minutes, allowing for switch-like effects on target protein structure and function. Previous mechanistic studies using small peptides derived from target substrates revealed Pin1 specifically binds to and displays enzymatic catalysis specificity for substrates containing a phosphorylated serine or threonine followed by a proline (pS/pT-P). Using a large substrate domain from the nuclear receptor peroxisome proliferator activated receptor gamma (PPARγ), we found that Pin1-catalyzed isomerization can occur at a noncanonical proline distal to a canonical pS/pT-P binding site. Our findings expand the understanding of Pin1-catalyzed enzymatic activities and target substrate functions.

## INTRODUCTION

Proline stands apart from other amino acids in its unconventional cyclic nature—an amino acid side chain that attaches to the peptide backbone at its amide nitrogen and imparts structural features which can play critical roles in the biological function of proteins. While most amino acid peptide bonds heavily bias *trans* conformations, the peptide bonds of proline can populate stable *cis* conformations (1, 2). Certain types of amino acids preceding a proline residue increase the propensity of adopting a *cis* conformation, including phosphorylated serine (pS) and threonine (pT), and aromatic residues, phenylalanine, tyrosine, and tryptophan (3, 4). Exchange, or isomerization, between *cis* and *trans* proline conformations in proteins is a relatively slow process that occurs on the order of seconds (5, 6). However, evolution has taken advantage of this dynamic quirk that alters protein function in what is commonly considered a switch-like mechanism (7–10). Proline *cis*-*trans* isomerization influences the activity of many proteins and plays central roles in a myriad of cellular processes including DNA damage response (11–13), cellular localization (14), ion channel activity (15), gene transcription (16, 17), and others (8, 18–21). Enzymes known as peptidyl prolyl *cis-trans* isomerases (PPIases) function as molecular timekeepers by catalyzing a noncovalent post-translational modification, the rate of exchange between *cis* and *trans* proline conformations, by several orders of magnitude (22, 23).

Peptidyl-prolyl *cis-trans* isomerase NIMA-interacting 1 (Pin1) is a multidomain hub PPIase that interacts with other proteins and initiates signaling cascades essential for cellular homeostasis (8, 24). Relative to other members of the PPIase superfamily, Pin1 uniquely contains a N-terminal WW domain (residues 1-39), which is connected to a C-terminal PPIase domain (residues 50-163) by a flexible linker. The WW domain directs Pin1 binding towards protein targets with consensus motifs containing a pS or pT N-terminally adjacent to a proline residue (pS/pT-P) (25–27). The PPIase domain, which enacts the catalytic *cis-trans* isomerase activity (28), was shown to preferentially catalyze isomerization of the peptide backbone between the pS/pT and proline residues (25, 29), though sequences with an acidic residue N-terminal to a proline were also reported to serve as a Pin1 enzymatic substrate (27).

Structures of Pin1 bound to phosphorylated peptides derived from target proteins determined by X-ray crystallography and NMR spectroscopy have detailed the molecular basis of substrate recognition. These studies have informed a model of interdomain allostery within Pin1 that occurs between the WW and PPIase domains, where target peptide binding to the WW domain primes the PPIase domain to operate with optimal catalytic capacity, as evidenced by target peptide-induced exchange between extended and compact Pin1 multidomain conformations (30–35). However, a more complete model of Pin1-target substrate interaction dynamics is lacking. Few studies have focused on Pin1-mediated recognition and enzymatic catalysis of substrates larger than peptides, such as entire protein domains that commonly include intrinsically disordered regions (IDRs) rich in clusters of post translational modifications. Studies using larger protein binding partners offer the opportunity to uncover more physiological binding and catalysis mechanisms and allosteric relationships within enzymatic substrates, which is especially relevant in the study of IDRs that contribute to complex protein signaling pathways.

Members of the nuclear receptor superfamily of ligand-regulated transcription factors are among the cellular substrates of Pin1 (36). Nuclear receptors contain a conserved modular domain architecture with a disordered N-terminal activation function-1 (AF-1) domain, a central DNA-binding domain, and a C-terminal ligand-binding domain (LBD) that contains the activation function-2 (AF-2) coregulator interaction surface and the ligand-binding pocket for orthosteric and synthetic ligands, including FDA-approved drugs (37). Two prior studies demonstrated Pin1 binds to and regulates the function of peroxisome proliferator activated receptor gamma (PPARγ), a lipid sensing nuclear receptor that regulates the expression of gene programs that influence insulin sensitization and the differentiation of mesenchymal stem cells into adipocytes (38). However, the studies suggested different Pin1 binding mechanisms to PPARγ, either to the disordered AF-1 domain (39) or folded LBD (40), and did not reveal a mechanism by which Pin1 accelerates *cis-trans* proline isomerization.

Obtaining a more comprehensive understanding of the structural basis for the binding and catalytic activity of Pin1 is key to understanding the role of proline *cis-trans* isomerization in the regulation of PPARγ-mediated transcription. In this study, we investigated the interaction between Pin1 and PPARγ using nuclear magnetic resonance (NMR) spectroscopy, which is ideally suited to map protein-protein interaction sites (41) and enzyme-catalyzed proline *cis-trans* isomerization (42). We found that Pin1 interacts with the isolated PPARγ AF-1 domain, but not the LBD. Phosphorylation of the AF-1 domain enhances Pin1 binding to the AF-1 through the interaction of the Pin1 WW domain with a pS-P motif within the AF-1. Moreover, Pin1 catalyzes *cis-trans* isomerization of a region within the AF-1 distal to the pS-P motif via a tethering mechanism containing an evolutionarily conserved tryptophan-proline (W-P) motif that is implicated in an interdomain interaction with the C-terminal ligand-binding domain (LBD) (43), suggesting a potential functional implication for Pin1-mediated *cis-trans* isomerization of the PPARγ AF-1 domain.

## RESULTS

### ERK2 phosphorylates human PPARγ AF-1 at two sites in vitro

Two well-characterized phosphorylation sites in PPARγ containing pS-P motifs are implicated by previous studies in binding to Pin1, including S112 in the AF-1 domain (in PPARγ2 isoform numbering, which corresponds to S84 in the shorter PPARγ1 isoform), which is phosphorylated by ERK2 (44), and S273 in the LBD, which is phosphorylated by Cdk5 (45) and ERK2 (46). We first sought to establish conditions for *in vitro* phosphorylation of these sites to test whether Pin1 interacts with the PPARγ2 AF-1 domain or LBD in a phosphorylation-dependent manner. Pin1 does interact with PPARγ2 AF-1, but not the LBD in unphosphorylated states (***SI Appendix*, Fig. S1**).

We performed *in vitro* phosphorylation of the PPARγ LBD using ERK2 and CDK5 but failed to detect phosphorylation of S273 in the LBD via phos-tag SDS-PAGE under the conditions tested (***SI Appendix*, Fig. S2**). However, *in vitro* phosphorylation of purified human PPARγ2 AF-1 domain by ERK2, followed by phos-tag SDS-PAGE, revealed two sequential phosphorylation events that occurs with different kinetics (***SI Appendix*, Fig. S2**). Mass spectrometry analysis of samples excised from the phos-tag SDS-PAGE revealed the faster ERK2-mediated phosphorylation occurs at S112 (pS112) and the slower phosphorylation occurs at T75 (pT75) (**Figure 1A**). We confirmed the time-dependent double phosphorylation via NMR spectroscopy with ^15^N-labeled AF-1 domain where ERK2 was added to the sample and a series of 2D [^1^H,^15^N]-HSQC NMR spectra were collected to monitor phosphorylation (**Figure 1B**). In the initial time points, the NMR time series data revealed disappearance of the NMR peak for S112 concomitant with the appearance pS112 in both *cis* and *trans* populations downfield (i.e., shifted to the left) from the unphosphorylated form. At the same time, the NMR peak for T75 decreased concomitant with population of pT75 NMR peaks downfield but at a much slower rate (tens of hours to days) compared to pS112 (minutes to hours), although no obvious *cis* pT75 population was observed.

**Figure 1.**
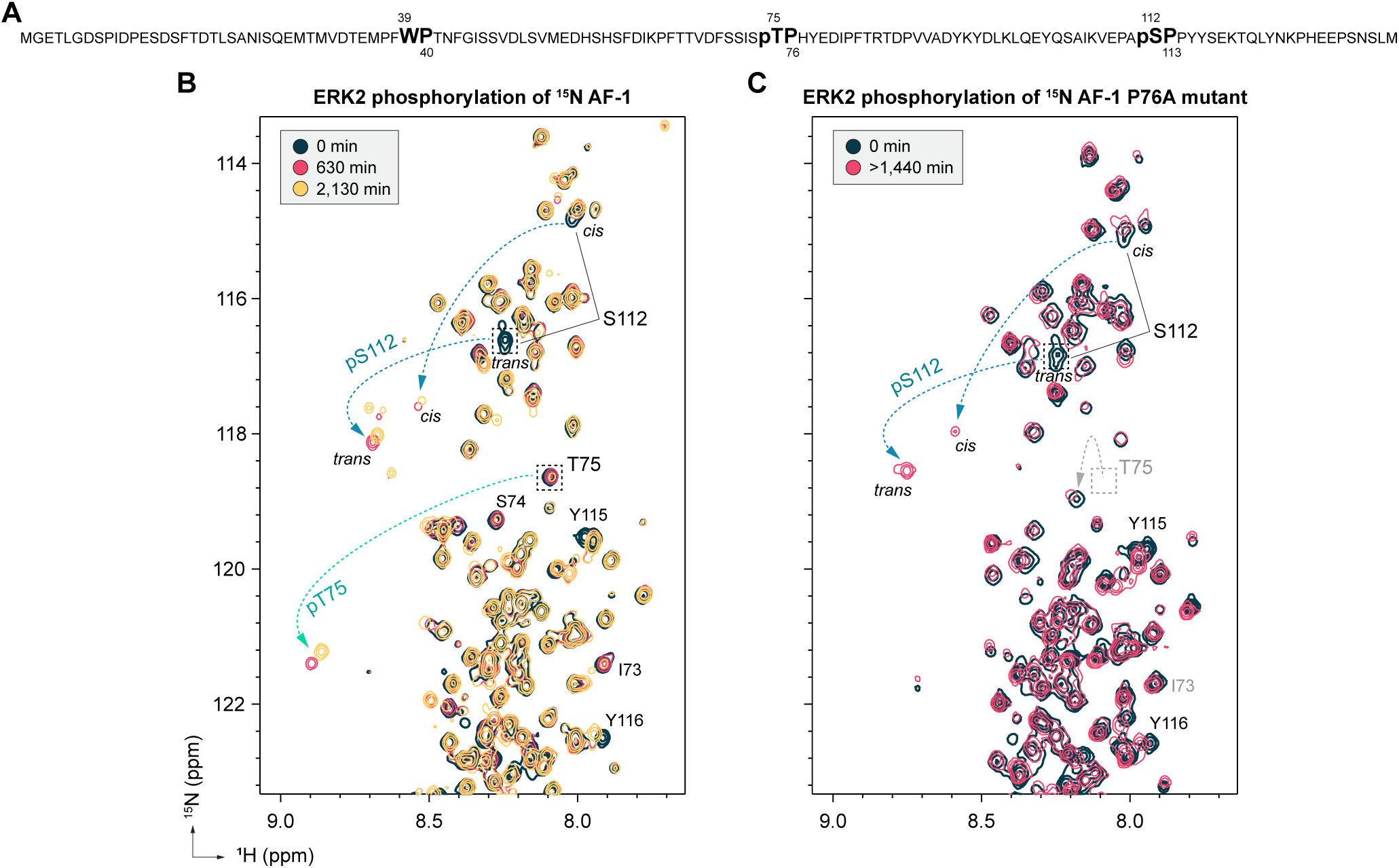
Two sites in the PPARγ AF-1 domain are phosphorylated by ERK2. (**A**) *In vitro* ERK2 phosphorylation sites within the AF-1 (pS112 and pT75) identified by mass spectrometry. A site that undergoes *cis-trans* isomerization that was previously identified (W39-P40) is also annotated. (**B**) ERK2 phosphorylation of ^15^N-labeled AF-1 followed by time-course 2D [^1^H,^15^N]-HSQC NMR confirms both phosphorylation sites (pS112 and pT75). (**C**) ERK2 phosphorylation of ^15^N-labeled AF-1 P76A mutant followed by time-course 2D [^1^H,^15^N]-HSQC NMR reveals only a single phosphorylation site (pS112). Gray dashed box and arrow indicate the shift of the peak corresponding to T75 which occurs due to mutation of the adjacent proline.

The putative T75 phosphorylation site is unique to human PPARγ2, as the corresponding residue in mouse PPARγ2 is an alanine residue (A75). To our knowledge, all previous studies that have focused on phosphorylation of PPARγ AF-1 domain used mouse receptor and mouse cell lines, and phosphorylation of T75 in human PPARγ has not been reported. Sequences of all peptides and protein constructs used in these studies (both Pin1 and PPARγ) are the respective human orthologs. To determine if human PPARγ2 may be phosphorylated at T75 in cells, we overexpressed full-length human PPARγ2 in human HEK293T cells and performed mass spectrometry on phosphoenriched peptides. Several sites in the AF-1 domain of PPARγ2 were phosphorylated, including several known (S26, S112) and previously unreported (S8, S51, S57, S72) sites; however, we did not detect phosphorylation of T75. It is possible T75 may be phosphorylated in other human cell types or tissues, or under other cellular conditions not tested such as cytokine signaling. We concluded our studies should account for binding contributions specific to phosphorylation at S112 as this rapid reaction (**Figure 1B**) is a previously validated modification that is known to enhance Pin1 binding to the PPARγ AF-1 (39), and because the physiological relevance/occurrence of phosphorylation at T75 remains ambiguous.

MAPK kinases including ERK2 specifically phosphorylate serine or threonine residues N-terminal adjacent to a proline residue (47, 48). We created an AF-1 domain construct where the proline C-terminal to T75 is mutated to alanine (P76A). *In vitro* phosphorylation of this AF-1 P76A mutant by ERK2 followed by 2D [^1^H,^15^N]-HSQC NMR (**Figure 1C**) only showed a single phosphorylation event that again resulted in the disappearance of the NMR peak for S112 concomitant with the appearance of downfield NMR peaks corresponding to pS112 in both *cis* and *trans* populations. Thus, the P76A mutant will allow the study of phosphorylation-dependent Pin1-mediated structure-function studies on human PPARγ AF-1 domain focusing on the pS112 site. The NMR peak corresponding to T75 (gray box, **Figure 1C**) is shifted relative to wild-type AF-1 because the P76 mutation changed the local chemical environment, but shows miniscule change in intensity relative to the unphosphorylated form and no peaks appear in the downfield region of the spectrum, which is characteristic of phosphorylated serine and threonine residues (49).

### AF-1 phosphorylation enhances binding to Pin1

Pin1 was previously shown to bind specifically to a peptide containing the PPARγ AF-1 pS112/P113 motif, but not in the unphosphorylated form (39). With a method established for ERK2-mediated phosphorylation of the AF-1 at a single site (pS112 via the P76A mutant) or two sites (pS112 + p75 via wild-type AF-1), we used 2D [^1^H,^15^N]-HSQC NMR analysis to structurally map where human Pin1 interacts with the AF-1 in phosphorylated and unphosphorylated states. Addition of Pin1 to unphosphorylated ^15^N-labeled AF-1 (**Figure 2A**) revealed subtle chemical shift perturbations (CSPs) in fast exchange on the NMR time scale suggestive of weak binding (5, 50) at select residues, particularly those C-terminal to the pT75 phosphorylation site, including E79, I81, T84, T86, and E87 (peaks denoted by blue labeling in **Figure 2A**). In contrast, Pin1 binding to phosphorylated ^15^N-labeled AF-1 (**Figure 2B**) or phosphorylated ^15^N-labeled P76A mutant (**Figure 2C**) caused more pronounced CSPs throughout the AF-1 with several noteworthy features. Residues near the pS112/P113 site that display *cis* and *trans* conformations, including E109, A111, and pS112 (peaks denoted by blue labeling in **Figures 2B and 2C**), consolidate to a single apparent *trans* conformation in the Pin1-bound state as the NMR peak corresponding to the *cis* conformation disappears. Furthermore, the pattern of CSPs that occurs for other residues throughout the AF-1 indicate that Pin1 interacts with multiple AF-1 regions or undergoes a conformational change upon binding to Pin1. Residues C-terminal to the pT75 and pS112 sites are generally affected on the intermediate exchange NMR time scale resulting in NMR peak line broadening, which could be suggestive of stronger binding compared to CSPs in fast exchange (5, 50). Moreover, a region of the AF-1 containing a PFWP motif (P37-F38-W39-P40; F38 and W39 denoted by green labeling in **Figures 2B and 2C**) that undergoes *cis-trans* isomerization (43) shows CSPs that occur on the fast exchange time scale, suggesting this region may bind more weakly to Pin1. Plotting CSP values calculated from data in **Figure 2B** on the top scoring AlphaFold 3 model of pAF-1 bound to Pin1 reveals the extent to which regions of AF-1 distal to the canonical pS112 binding site of Pin1 engage with the isomerase (**Figure 2D**). In agreement with previously reported models (39) and experimental data shown further in this results section, AlphaFold3 structures of pAF-1 bound to Pin1 model pS112 in close proximity to the WW domain.

**Figure 2.**
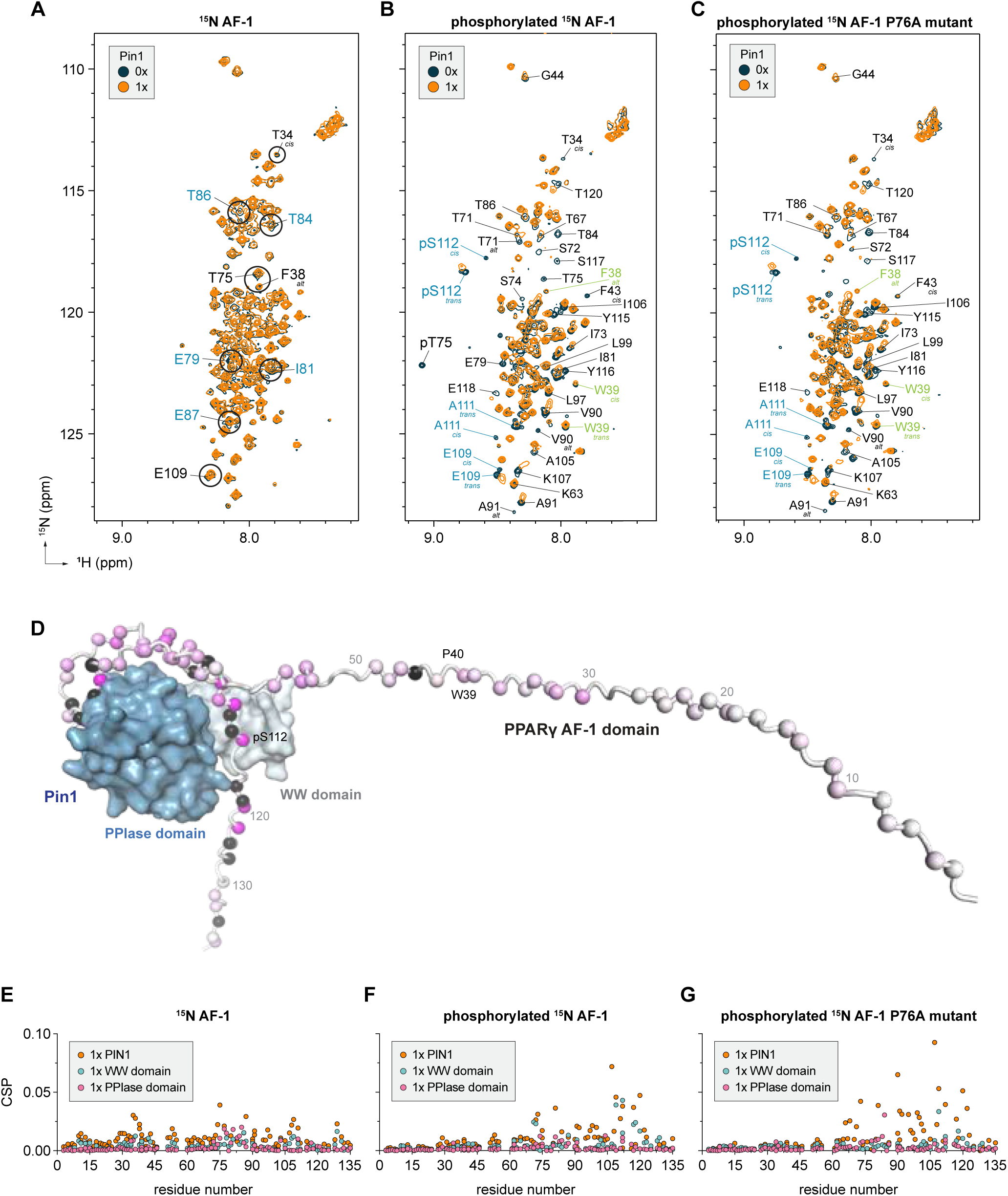
Pin1 binding to the AF-1 is enhanced by ERK2 phosphorylation. Overlays of 2D [^1^H,^15^N]-HSQC NMR spectra of (**A**) ^15^N-labeled AF-1, (**B**) ^15^N-labeled pAF-1, and (**C**) ^15^N-labeled pAF-1 P76A mutant without or with 1 molar equivalent of human Pin1 reveal larger chemical shift perturbations in the phosphorylated state indicating stronger binding of Pin1 to phosphorylated AF-1. Peaks belonging to regions of particular interest noted by blue and green labeling are discussed further in the results. (**D**) AlphaFold3 model of Pin1 interaction with pAF-1 containing phosphorylated S112 (pS112) and T75 (pT75) residues with NMR CSPs from ^15^N-labeled pAF-1 ± 1x Pin1 (**2B**) shows the Pin1-binding epitope on pAF-1 extends to regions beyond the pS112 region. Spheres represent residue alpha carbons colored by CSP magnitude (white (CSP=0) to magenta (CSP=0.03)) or noting peaks that disappear (black). Chemical shift perturbation (CSP) plots comparing addition of Pin1 full-length, WW domain, or PPIase domain at 1 molar equivalent to (**E**) ^15^N-labeled AF-1, (**F**) ^15^N-labeled pAF-1, and (**G**) ^15^N-labeled pAF-1 P76A mutant reveal that each Pin1 domain binds weakly to AF-1, but binding of full-length Pin1 elicits larger CSPs in targeted regions of pAF-1 and pAF-1 P76A mutant.

### Mapping Pin1 domain binding contributions to PPARγ AF-1

The 2D NMR structural footprinting data indicate that multiple regions of the PPARγ AF-1 comprise the Pin1-interaction surface. To distinguish the binding contributions of the Pin1 WW domain (residues 1–39) and PPIase domain (residues 42–163), we compared 2D [^1^H,^15^N]-HSQC NMR spectra of ^15^N-labeled AF-1 in unphosphorylated and phosphorylated states along with the isolated domains of Pin1.

Addition of the WW domain to unphosphorylated ^15^N-labeled AF-1 resulted in negligible CSPs (**Figures 2E and *SI Appendix*, Fig. S3A**) but caused notable CSPs for phosphorylated ^15^N-labeled AF-1 (**Figures 2F and *SI Appendix*, Fig. S3B**) and phosphorylated ^15^N-labeled AF-1 P76A mutant (**Figures 2G and *SI Appendix*, Fig. S3C**). In the phosphorylated ^15^N-labeled AF-1 analysis, residues with larger CSPs map to the pS112/P113 and pT75/P76 phosphorylation sites, whereas only the pS112/P113 site is affected in the phosphorylated ^15^N-labeled AF-1 P76A mutant analysis. These data are consistent with published studies indicating that the Pin1 WW domain specifically recognizes and binds to pS/pT-P motifs (25–27).

Addition of the PPIase domain to unphosphorylated ^15^N-labeled AF-1 (**Figures 2E and *SI Appendix*, Fig. S3D**) resulted in minor but discernible CSPs for select residues, including residues within and near the PFWP motif (W39 *cis* and *trans* peaks, T34 *cis* peak), within and near the T75 phosphorylation motif (S74, T75), as well as a nearby downstream site (E79, I81, T84, R85, T86). These residues are also affected when the PPIase domain is added to phosphorylated ^15^N-labeled AF-1 (**Figures 2F and *SI Appendix*, Fig. S3E**) and phosphorylated ^15^N-labeled AF-1 P76A mutant (**Figures 2G and *SI Appendix*, Fig. S3F**); however, the pS112 *cis* and *trans* peaks and pT75 peak also shows notable CSPs. These data indicate the Pin1 PPIase domain may recognize or interact with multiple regions of the PPARγ AF-1.

Direct comparison of CSPs observed by addition of individual Pin1 domains versus full length Pin1 to samples of ^15^N-labeled AF-1 in both phosphorylated and unphosphorylated states provided additional insight into the Pin1 binding mechanism. Full-length Pin1 binding to unphosphorylated ^15^N-labeled AF-1 reveals larger CSPs compared to CSPs observed in the presence of the WW domain or PPIase domain alone (**Figure 2E**). Furthermore, CSPs caused upon binding full-length Pin1 appear at distinct AF-1 regions including near the PFWP motif (residues 35-39), near the T75 phosphorylation site (residues 75-90), near the S112 phosphorylation site (residues 105-110), and a region near the C-terminus of the AF-1 among other regions. In contrast, AF-1 phosphorylation appears to fine-tune the interaction. Binding of full-length Pin1 and, to a lesser degree, the WW domain to phosphorylated ^15^N-labeled AF-1 (**Figure 2F**) or phosphorylated ^15^N-labeled AF-1 P76A mutant (**Figure 2G**) shows larger CSPs compared to unphosphorylated AF-1 that are localized to the C-terminal region of the AF-1 with the largest CSPs occurring around the S112 phosphorylation site. Moreover, CSPs caused by binding full-length Pin1 are much larger than that of either domain alone.

### Mapping PPARγ AF-1 binding surface on Pin1

We next used 2D [^1^H,^15^N]-TROSY HSQC NMR analysis to structurally map where the AF-1 interacts with Pin1 in phosphorylated and unphosphorylated AF-1 states. Addition of unphosphorylated AF-1 to ^15^N-labeled Pin1 (**Figure 3A**) resulted in CSPs for select residues within the WW domain and PPIase domain. However, addition of phosphorylated AF-1 to ^15^N-labeled Pin1 (**Figure 3B**) resulted in much larger CSPs, indicating the interaction is more robust than with unphosphorylated AF-1. CSP analysis revealed that AF-1 phosphorylation significantly increases interaction with the WW domain (**Figure 3C**). Moreover, AF-1 phosphorylation results in larger CSPs for residues within the PPIase domain comprising the regions within and near the catalytic loop (residues 70-75) and catalytic active site that includes residues H59, C113, S154, and H157. Furthermore, peak intensity analysis reveals a larger decrease in NMR peak intensities for the Pin1 WW domain and PPIase domain upon interacting with phosphorylated AF-1 compared to unphosphorylated AF-1 (**Figure 3D**), further indicating a more robust interaction.

**Figure 3.**
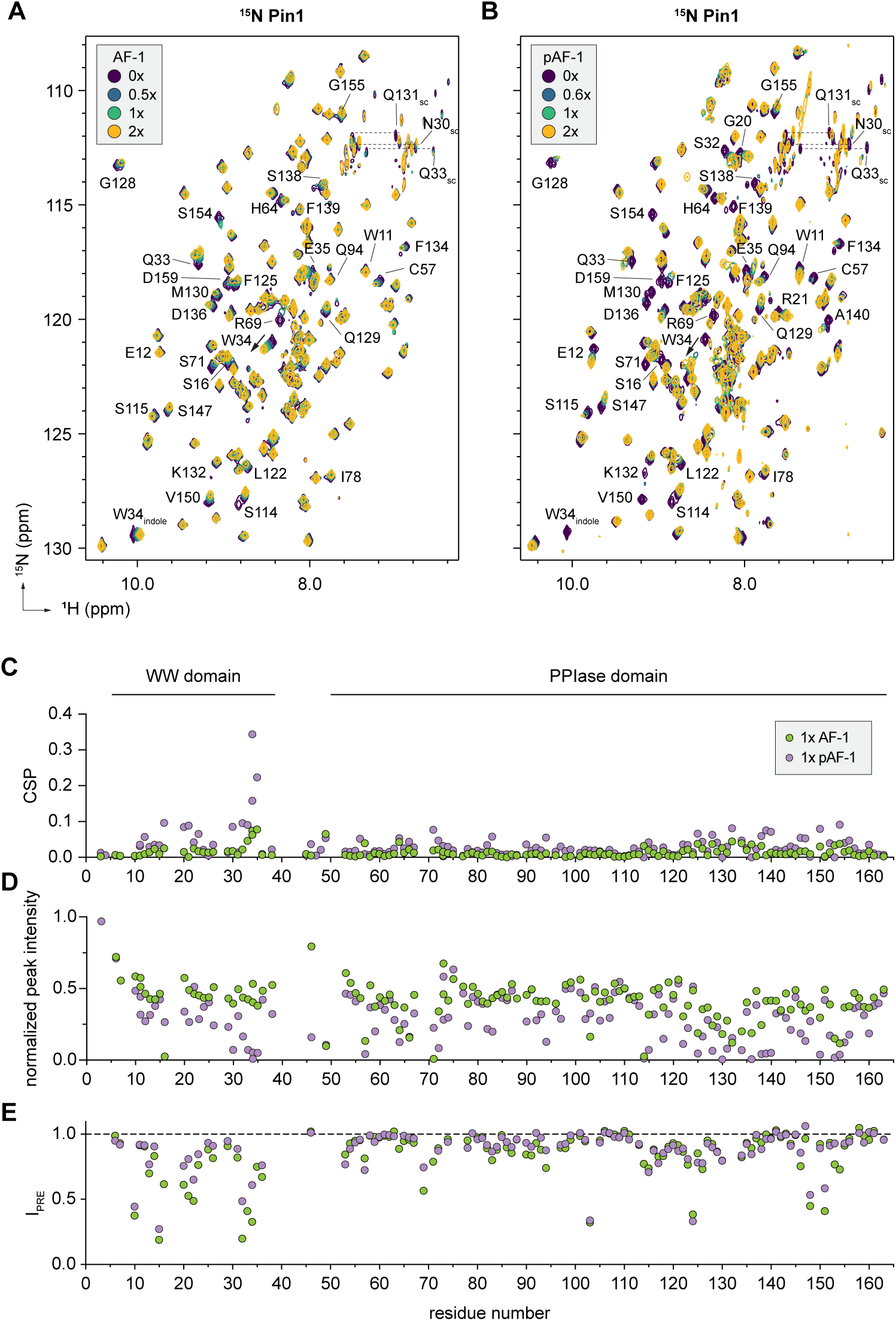
AF-1 phosphorylation drives binding to the Pin1 WW domain. Overlays of 2D [^1^H,^15^N]-TROSY HSQC NMR spectra of ^15^N-labeled Pin1 without or with 0.5, 1, or 2 molar equivalents of (**A**) AF-1 or (**B**) pAF-1. (**C**) Chemical shift perturbation (CSP) and (**D**) normalized peak intensity plots comparing addition of AF-1 or pAF-1 at 1 molar equivalent to ^15^N-labeled Pin1. (**E**) Paramagnetic relaxation enhancement (PRE) NMR using MTSSL-labeled pAF-1 D33C+T75A mutant. The data show that phosphorylated AF-1 binding to Pin1 displays larger CSPs and thus a stronger interaction compared to unphosphorylated AF-1, driven in large part by interactions with the Pin1 WW domain.

We used paramagnetic relaxation enhancement (PRE) NMR to more specifically detect the AF-1 PFWP motif-binding surface in Pin1. For these PRE NMR studies, we used an AF-1 double mutant protein (D33C/T75A) containing a D33C mutation to attach the MTSSL spin probe, which we used previously to show the PFWP motif interacts with the PPARγ LBD (43), as well as a T75A mutation to limit ERK2 phosphorylation to S112. Addition of MTSSL-labeled AF-1 D33C/T75A mutant protein to ^15^N-labeled Pin1 revealed PRE NMR effects within the Pin1 WW domain, linker, and PPIase domain (**Figure 3E, *SI Appendix*, Fig. S4**). MTSSL-labeled phosphorylated AF-1 D33C/T75A mutant protein revealed a similar profile. However, PRE effects within the WW domain caused by the phosphorylated AF-1 were attenuated or not as robust as the unphosphorylated form. One interpretation of these data is that phosphorylation of S112 affords higher affinity binding of the pS112-P113 region to the WW domain, which would reduce the chance of transient or robust interactions between the PFWP motif region to the WW domain.

To explore the specific Pin1-binding regions of the AF-1 surfaces identified in the differential NMR analysis of ^15^N AF-1, we performed 2D [^1^H,^15^N]-TROSY HSQC NMR titrations where ^15^N-labeled Pin1 was incubated with increasing concentrations of peptides derived from three AF-1 regions including the two phosphorylation sites without (S112 and T75) and with phosphorylation (pS112 and pT75), as well as the PFWP motif (**Figure 4A-F**) and performed CSP analysis (**Figure 4G**). Binding of the unphosphorylated peptides caused subtle but discernable CSPs for residues in the Pin1 WW and PPIase domains, suggestive of a weak interaction compared to the phosphorylated peptides. Of the unphosphorylated peptides, the peptide comprising the AF-1 region including T75 showed the largest effects. The PFWP-containing region showed more moderate effects as did the S112 region. In contrast, binding of the phosphorylated pS112 and pT75 peptides revealed larger CSPs focused primarily within the WW domain.

**Figure 4.**
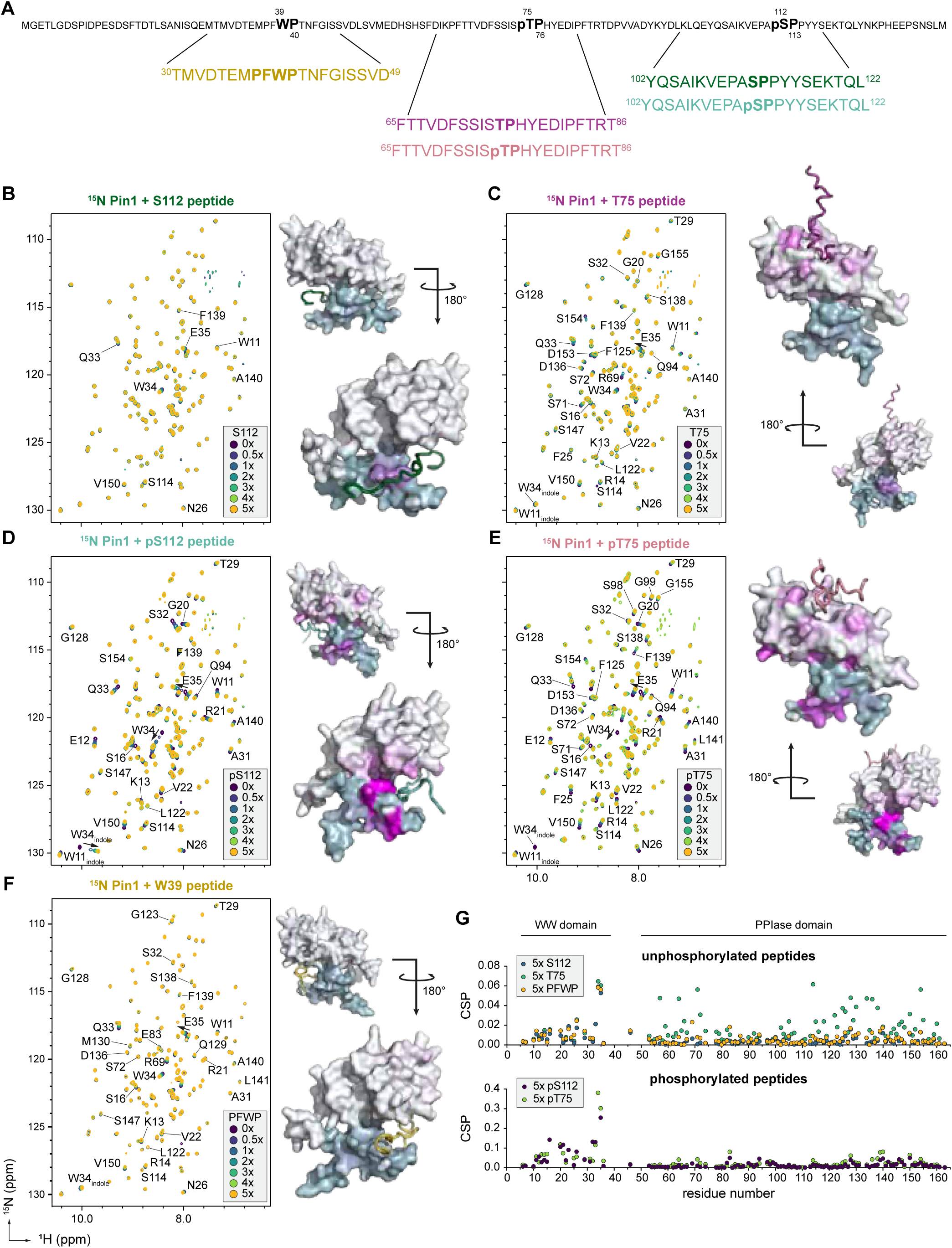
AF-1 peptides delineating contributions of specific AF-1 regions in binding to Pin1. (**A**) Sequence and relative locations of peptides corresponding to AF-1 regions that include the S112 phosphorylation site with unphosphorylated S112 (green) or phosphorylated S112 (teal); T75 phosphorylation site with unphosphorylated T75 (purple) or phosphorylated T75 (coral); and the PFWP motif (mustard). Overlays of 2D [^1^H,^15^N]-TROSY HSQC NMR spectra of ^15^N-labeled Pin1 with (**B**) S112, (**C**) T75, (**D**) pS112, (**E**) pT75, and (**F**) W39 peptides titrated up to 5 molar equivalents. Shown alongside spectra are AlphaFold3 models of Pin1 colored by magnitude of CSP (pale blue/white (CSP=0) to magenta (CSP=0.1)) elicited by 2X titration of respective peptide shown in complex with Pin1. WW domain is colored pale blue and PPIase domain is white. Peptides in models are colored corresponding to their identity as indicated in **5A**. (**G**) Chemical shift perturbation (CSP) plots comparing addition of unphosphorylated or phosphorylated peptides at 5 molar equivalents to ^15^N-labeled Pin1. CSP profiles reveal Pin1 interaction patterns unique to each AF-1 peptide, with phosphorylated AF-1 peptides driving enhanced interaction to the WW domain.

### Affinity calculations between Pin1 and the AF-1

We used NMR TITAN lineshape analysis software (51, 52) to determine domain specific affinity interactions (**Table 1**). Addition of phosphorylated AF-1 to ^15^N Pin1 (***SI Appendix*, Fig. S5, Dataset S1**) resulted in a fitted WW domain specific *K_D_* of 4.44 ± 0.23 µM, in good agreement with the reciprocal measurement using titrations of unlabeled Pin1 to ^15^N-labeled pAF-1 (***SI Appendix*, Fig. S6, Dataset S2**) resulting in a fitted *K_D_* specific to the region around pS112 of 4.08 ± 0.93 µM. Our attempts to use NMR TITAN to obtain *K_D_* values from titration data using unphosphorylated AF-1 failed to reliably reproduce fits, likely due to several reasons: multiple weak Pin1 binding sites within the AF-1 (S112 site, T75 site, W39-P40 site), multiple weak AF-1 binding sites on Pin1 (PPIase domain, WW domain, hinge region between these domains), and the fact that more than one region of the AF-1 that interacts with Pin1 undergoes *cis-trans* isomerization. However, we note that our attempted fittings indicate Pin1 binding to unphosphorylated AF-1 occurs with an affinity at least/more than an order of magnitude weaker than pAF-1.

**Table 1.** NMR TITAN fitted affinity measurements for Pin1/pAF-1 binding interaction.

| Sample (residues included) | $K_D$ ($\mu\text{M}$; ± std. err.) | $k_{\text{off}}$ ($\text{s}^{-1}$; ± std. err.) |
| --- | --- | --- |
| **$^{15}\text{N}$-labeled pAF-1 + Pin1** |  |  |
| $^{15}\text{N}$ pAF-1 (200 $\mu\text{M}$); Pin1 (0X, 0.2X, 0.4X, 0.6X, 0.8X, 1X) | | |
| E109, pS112 | $4.08 \pm 0.93$ | $659 \pm 142$ |
| **$^{15}\text{N}$-labeled Pin1 + pAF-1** |  |  |
| $^{15}\text{N}$ Pin1 (200 $\mu\text{M}$); pAF-1 (0X, 0.2X, 0.4X, 0.6X, 0.8X, 1X) | | |
| WW domain (W11, E12, R14, M15, G20, R21, V22, F25, N26, A31) | $4.44 \pm 0.23$ | $1433 \pm 71.4$ |

### Pin1 catalyzes cis-trans isomerization of a noncanonical WP motif in the AF-1

The AF-1 peptide and AF-1 domain NMR titrations into ^15^N-labeled Pin1 suggest a model whereby phosphorylation drives interaction of the pS/pT motif to the WW domain. However, additional interactions occur between the Pin1 PPIase domain and hinge region to other pAF-1 regions (**Figure 2D**) including the W39-P40 motif that undergoes *cis-trans* isomerization (43).

Orienting the interacting regions of Pin1 and AF-1 through CSP and PRE NMR analyses led us to hypothesize that Pin1 may be able to accelerate *cis-trans* isomerization of the noncanonical WP dipeptide motif in the AF-1. We previously used 3D C(CCO)NH-TOCSY NMR and 2D [^1^H,^15^N]-ZZ exchange NMR, which detects conformational exchange that occurs on slow time scales (10–5,000 ms) including proline *cis-trans* isomerization processes (5), to show that the WP dipeptide sequence within the PFWP motif undergoes *cis-trans* isomerization (43). In that study, ZZ exchange NMR could detect the slow conformational exchange of the WP motif at elevated temperatures (>35 °C) but not at room temperature (25 °C), indicating the exchange process is on the order of seconds or longer (k_ex_ ≈ 0.2-100/s; τ_ex_ ≈ 10-5,000 ms) (5). Consistent with these prior findings, 2D [^1^H,^15^N]-ZZ exchange NMR of ^15^N-labeled AF-1 in the unphosphorylated and phosphorylated states at 25 °C revealed no apparent exchange peaks using an exchange delay up to 1.5 seconds (**Figure 5A**). Strikingly, addition of equimolar Pin1 (**Figure 5B**) and sub-stoichiometric catalytic concentrations of Pin1 (**Figure 5C**) accelerated *cis-trans* isomerization of the WP dipeptide motif in both the unphosphorylated and phosphorylated states at 25 °C, evident by the appearance of ZZ exchange correlation NMR cross peaks.

**Figure 5.**
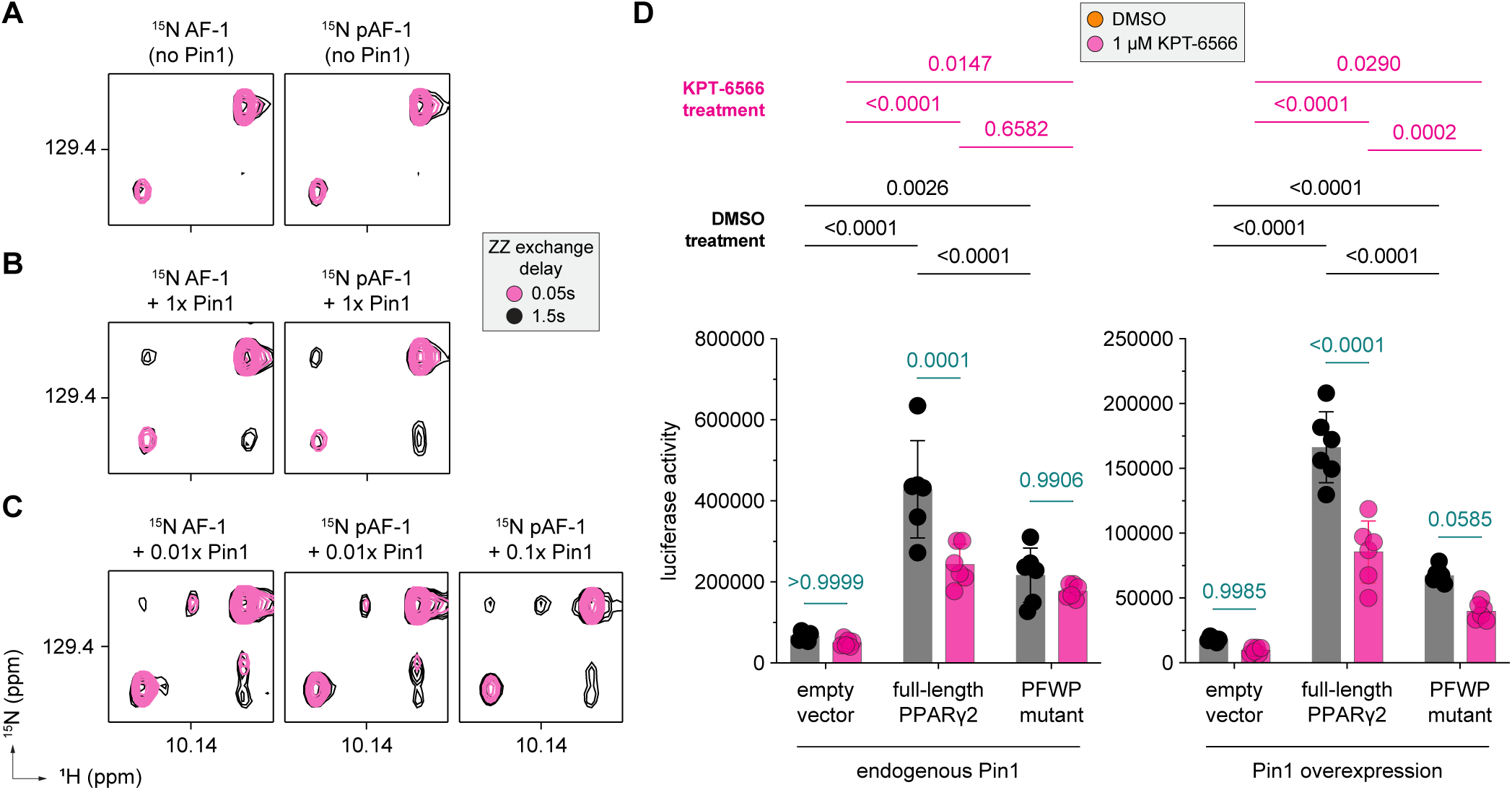
Pin1 accelerated *cis-trans* isomerization of WP motif influences PPARγ transcription. Overlays of 2D [^1^H,^15^N]-ZZ exchange NMR spectra of ^15^N-labeled AF-1 or pAF-1, zoomed into the W39 indole NMR peaks, in the absence (**A**) or presence of equimolar (**B**) or subequimolar catalytic (**C**) equivalents of Pin1 demonstrate that Pin1 accelerates *cis-trans* isomerization of the W39-P40 dipeptide motif. (**D**) Cellular luciferase transcriptional reporter assay in HEK 293T cells using a 3xPPRE-luciferase reporter plasmid with overexpression of PPARγ or a PFWP motif mutant (PFWP to AAAA) in the absence or presence of 1 µM KPT-6566, a covalent inhibitor of the Pin1 PPIase domain, with or without overexpression of Pin1 (n=6; mean ± s.d.). Multiplicity adjusted P values are indicated above chart.

To determine if Pin1-catalyzed acceleration of the WP motif influences transcription, we performed a cellular transcriptional reporter assay (**Figure 5D**). Cells were transfected with full-length PPARγ along with a second plasmid containing three copies of the PPAR DNA-binding response element sequence (3xPPRE) upstream of luciferase gene. Experiments were performed in the presence of endogenous Pin1 and with co-transfection of a Pin1 expression plasmid — with and without a pharmacological Pin1 inhibitor, KPT-6566, which covalently binds to the Pin1 PPIase domain and blocks its enzymatic activity (53). Transfection of full-length PPARγ increased transcriptional activity, which was decreased upon KPT-6566 treatment, indicating that Pin1-catalyzed *cis-trans* isomerization activates PPARγ-mediated transcription and endogenous levels of Pin1 present within cells (54), which we verified using RT-qPCR (PIN1 Ct value ∼24.2 vs. GAPDH Ct value ∼18.5), are sufficient to produce the response. Although endogenous PIN1 is abundant as evidenced by Ct values relative to GAPDH from RT-qPCR analysis, trends mirrored across conditions with and without Pin1 overexpression give credence to the notion that catalytic activity and not stoichiometric binding account for the mechanism by which Pin1 mediates PPARγ transcriptional activity. Transfection of a mutant PPARγ variant where the PFWP motif was mutated to four consecutive alanine residues (PFWP to AAAA) also increased activity of the luciferase reporter, but to a lower degree compared to wild-type PPARγ. However, KPT-6566 treatment had no significant effect on the activity of the PFWP mutant, suggesting that Pin1-catalyzed *cis-trans* isomerization of the WP motif may be a primary mechanism by which Pin1 influences catalysis-induced effects on PPARγ-mediated transcription. Similar trends were observed when Pin1 was overexpressed in cells, though one difference is KPT-6566 decreased activity of the PFWP mutant indicating overexpression of Pin1 increases sensitivity of the Pin1 inhibitor treatment.

## DISCUSSION

Current thinking in the field suggests that Pin1 selectively accelerates pS/pT-P motifs undergoing *cis-trans* isomerization (33, 55, 56). Several mechanistic models have been proposed for Pin1-mediated catalysis of pS/pT-P motifs within a single target, including a catalysis first mechanism where the PPIase domain first catalyzes the *cis*-to-*trans* isomerization followed by binding of the WW domain to the pS/pT-P motif in the *trans* conformation. The sequential model proposes the WW domain first binds to a pS/pT-P motif in the *trans* conformation, tethering the PPIase domain to the target and enabling catalysis of other substrate pS/pT-P motifs. Another related model is the simultaneous or bivalent binding model, where the WW and PPIase domains bind to low affinity target pS/pT-P motifs at the same time (57). These binding and catalysis models were developed from studies that used small peptide fragments from substrate targets. In contrast, our findings here using a large IDR substrate domain indicate the interaction between the PPARγ AF-1 domain and Pin1, and enzymatic catalysis of a noncanonical substrate motif (**Figure 6**), may combine features of all three mechanistic models.

**Figure 6.**
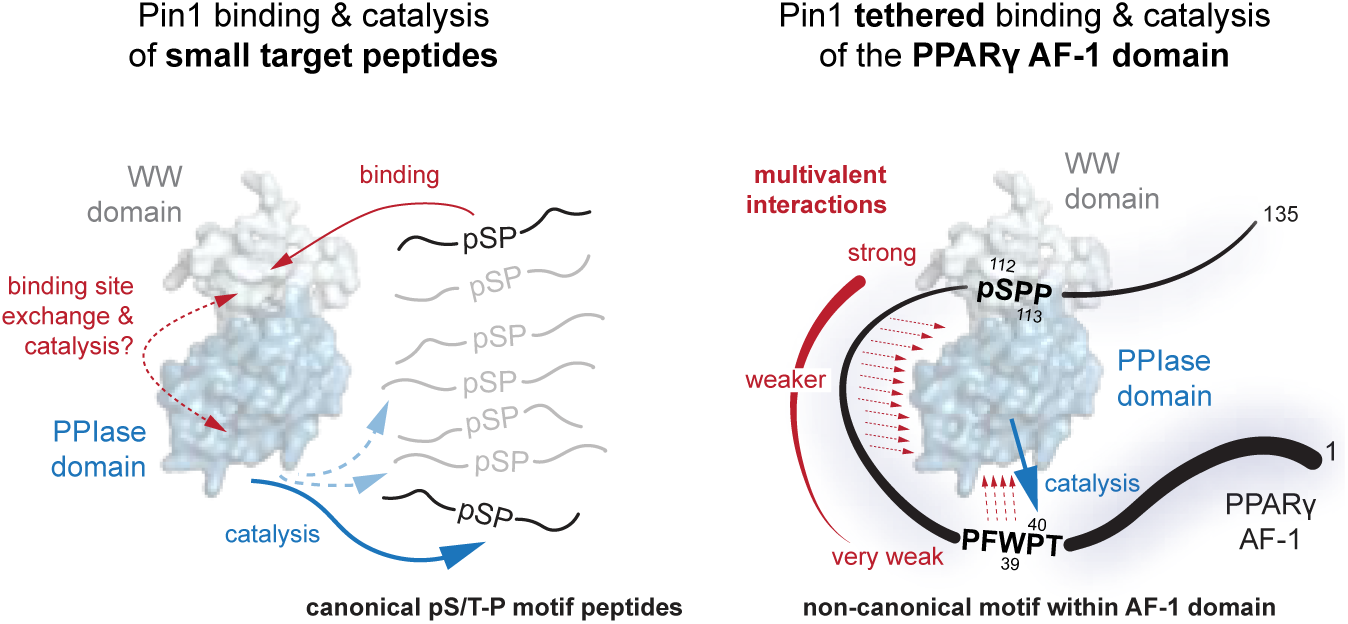
Approaches to studying Pin1 binding and catalysis. Previous structural studies of Pin1 isomerization depended heavily on the use of short peptide sequences (∼20 amino acids). In addition to short peptides, our studies using a larger protein domain with multiple potential canonical and noncanonical Pin1 sites uncovered a multivalent binding interaction that catalyzes a noncanonical W-P motif with functional cellular relevance.

In the Pin1-bound ^15^N-labeled pAF-1 NMR spectrum, an NMR peak is observed for pS112 in the *trans* conformation but not in the *cis* conformation. It is possible that Pin1 first catalyzed the *cis*-to-*trans* isomerization of pS112, which produced the Pin1-bound AF-1 NMR spectrum where only the pS112 *trans* conformation is observed. However, another interpretation of these data is that the Pin1 WW domain selectively binds to the pS112 *trans* conformation, and during sample preparation and NMR data collection the *cis* conformation natively exchanged to the *trans* conformation, the latter of which could then bind to Pin1 WW domain, eventually resulting in the loss of the pS112 *cis* conformation.

Our NMR CSP data shows that AF-1 phosphorylation drives binding of the Pin1 WW domain to the pS112-P113 site. While our studies demonstrate phosphorylation occurs at another canonical Pin1 binding motif, p75-P76, similar CSP profiles of doubly phosphorylated pAF-1 and a singly phosphorylated P76A mutant suggest pS112 alone is a potent driver of affinity towards Pin1. As titrations of T75 peptide shows, the region of AF-1 around the T75 phosphorylation site maintains affinity for Pin1 independent of phosphorylation of AF-1. These observations underscore the value of using extended protein domains in the study of Pin1 multivalent interactions and indicate a cooperative, allosteric mechanism may underlie the interaction between Pin1 and pAF-1 where the interaction involves multiple surfaces on each protein.

The nature of Pin1 as a hub protein which interacts with scores of other proteins belonging to classes including kinases, mitotic regulars, and transcription factors lends itself to the idea that Pin1 domains must retain some degree of substrate promiscuity. Complementarily, the disordered nature of PPARγ AF-1 imparts a large degree of conformational flexibility also permitting interaction with numerous binding partners, a common feature among similar regulatory IDRs. Indeed, others have gone as far as to describe the nature of IDP binding interactions as “fuzzy” (58–60). Our data here from CSP and PRE NMR, which are robust methods to detect weak or transient interactions (61), show that the AF-1 PFWP motif binds to multiple surfaces on Pin1 including WW and PPIase domains. Despite the apparent weak affinity of the AF-1 PFWP motif for Pin1, this motif—a noncanonical site that does not conform to the classical pS/pT-P target identity—undergoes Pin1-catalyzed *cis-trans* isomerization. The tethering of a binding partner such as AF-1 via a pS/pT-P motif to the WW domain may provide a molecular basis by which Pin1 can exert *cis-trans* catalysis of a noncanonical motif via its PPIase domain. Our cellular studies reveal there is a functional consequence of Pin1-catalyzed *cis-trans* isomerization of this noncanonical site, as the PFWP to AAAA mutant shows reduced transcriptional activity and significantly decreased sensitivity to a covalent Pin1 PPIase inhibitor. Our data support a modified tethering model that challenges the current Pin1 catalytic mechanism paradigm in a few important ways. While the interaction of the AF-1 pS/pT-P motifs to Pin1 via the WW domain are well aligned with current models of Pin1 binding, the ability of Pin1 to catalyze *cis-trans* isomerization of a noncanonical site in a target substrate and the detailing of multiple substrate regions interacting with both Pin1 domains extends the current models of Pin1-mediated enzymatic catalysis.

## MATERIALS AND METHODS

### Reagents

Activated recombinant human ERK2 (ab155812) and CDK5/p25 (ab60761) proteins for *in vitro* kinase reactions were purchased from Abcam. N-terminal acetylated peptides synthesized by Lifetein included (human PPARγ2 isoform numbering): S112 (^102^YQSAIKVEPASPPYYSEKTQL^122^), pS112 (^102^YQSAIKVEPApSPPYYSEKTQL^122^), T75 (^65^FTTVDFSSISTPHYEDIPFTRT^86^), pT75 (^65^FTTVDFSSISpTPHYEDIPFTRT^86^), and PFWP (^30^TMVDTEMPFWPTNFGISSVD ^49^). Pin1 inhibitor KPT-6566 (37308, CAS# 881487-77-0) was purchased from Cayman Chemicals and dissolved in DMSO-d_6_ before addition to NMR samples or use in reporter assays. MTSSL PRE NMR spin label (16463, CAS# 81213-52-7) was purchased from Cayman Chemicals.

### Protein expression and purification

Wild-type and mutant human PPARγ (isoform 2) activation function 1 (AF-1) protein (residues 1-136) was expressed from a pET45 plasmid vector as a TEV-cleavable N-terminal 6xHis fusion protein in *Escherichia coli* BL21(DE3) cells using terrific broth (TB) or M9 minimal media for expression of isotopically labeled (^15^N-labeled or ^15^N,^13^C-labeled) protein. Human Pin1 PPIase domain (42-163) was expressed from a pET24 plasmid vector (Twist biosciences) as a 3C-cleavable N-terminal 6xHis fusion proteins. Full-length human Pin1 and the Pin1 WW domain (1-39) protein were expressed from a pMCSG7 plasmid vector, purchased through Addgene (#40773) as TEV-cleavable N-terminal 6xHis fusion proteins. PPARγ ligand binding domain (residues 231-505) was expressed from a pET46 Ek/LIC plasmid (Novagen) as a TEV-cleavable N-terminal 6xHis fusion protein. Cells transformed with plasmid were initially grown at 37°C and 200 rpm to an OD_600nm_ of 1.2 (TB media) or 0.8 (M9 media), then supplemented with 1 mM isopropyl β-D-thiogalactoside (IPTG) for induction of protein expression at 18 °C for 16 h (Pin1 proteins) or at 37 °C for 4 h (AF-1 proteins). Cells were harvested and lysed by sonication after resuspension in lysis buffer (40 mM phosphate pH 7.4, 500 mM KCl, 0.5 mM EDTA, 15 mM imidazole), the lysate clarified by centrifugation at 20,000xg, filtered, and purified by Ni-NTA affinity chromatography (elution buffer: 40 mM phosphate pH 7.4, 500 mM KCl, 0.5 mM EDTA, 500 mM imidazole). TEV or 3C proteases were used to cleave 6xHis tags from proteins during overnight dialysis in imidazole free lysis buffer at 4 °C. Contaminant and tag removal was accomplished by an additional Ni-NTA affinity chromatography step, whereby cleaved protein was gathered from the wash phase of purification and further purified using size exclusion chromatography with an S75 column (GE healthcare) into NMR Buffer (20 mM phosphate pH 7.4, 50 mM KCl, 0.5 mM EDTA) for long term storage if protein samples were sufficiently pure. At this stage, AF-1 proteins were heated in a hot water bath at 80 °C for 15 minutes to denature any additional contaminating proteins and centrifuged at 4000 rpm for 15 minutes before concentration and purification by anion exchange chromatography through a HiTrap Q HP column (GE healthcare) using a high salt buffer to elute protein from the column (20 mM phosphate pH 7.4, 1M KCl, 0.5 mM EDTA). Pin1 proteins were purified by a final anion exchange chromatography as seen fit.

### In vitro phosphorylation

*In vitro* phosphorylation reactions were conducted in NMR Buffer supplemented with 10 mM MgCl_2_, 1 mM ATP, and Pierce Protease Inhibitor (ThermoFisher Scientific A32955). 1:3000 ERK2:AF-1 was used in all phosphorylation experiments. “Stopping” phosphorylation was achieved by heating reactions at 80 °C for 15 minutes to denature ERK2. For protein used in NMR experiments, samples were then dialyzed overnight in NMR Buffer to remove excess ATP and MgCl_2_. *In vitro* phosphorylation reactions were monitored using Super Sep Phos-tag precast gels (195-17991) or gels cast in house with Phos-tag acrylamide (AAL-107) from Fujifilm Wako Chemicals. In select experiments, bands of interest were excised from gels and subject to proteomic analysis to identify sites of phosphorylation. ^15^N or ^15^N,^13^C labeled proteins whose reaction progressions were monitored by NMR were conducted under identical conditions. ERK2 kinase was added after the collection of a completely unphosphorylated state spectrum of the protein.

### Mass spectrometry

To identify phosphorylation sites on proteins from bands excised from Super Sep Phos-tag gels, samples were digested and a 1-hour LC-MS/MS run was performed. The primary sequence (AF-1; residues 1-135 of PPARγ2 WT or S112A mutant) was appended to 4172 proteins from *E. coli* BL21 DE3 strain downloaded from Uniprot, for searching for peptide sequences. A common contaminant protein fasta file was also used in the search. Phosphorylation was considered on S, T, and Y residues and the location probability node was also used in the search. On bands corresponding to pS112 as indicated in ***SI Appendix*, Fig. S2B,** 68% protein sequence coverage was achieved with detection of phosphorylation at S112 alone using a probability threshold of 99%. On bands corresponding to pT75 + pS112 as indicated in ***SI Appendix*, Fig. S2B**, 68% protein sequence coverage was achieved with detection of phosphorylation at T75 and S112 using a probability threshold of 99%. More details and experimental reporting can be found on the MassIVE database (dataset MSV000096300).

### NMR spectroscopy

NMR experiments were collected at 298K on Bruker 600, 700, and 900 MHz NMR equipped with cryoprobes. 20 mM phosphate pH 7.4, 50 mM KCl, 0.5 mM EDTA buffer conditions were used for all samples. All samples contained 10% D_2_O for NMR instrument locking, except for the ^15^N AF-1 + Pin1 titration series that contained 5% DMSO-d_6_. NMR samples contained 200 µM ^15^N-labeled protein, except for samples prepared for 3D NMR (450+ µM) to validate NMR assignments via ^13^C,^15^N-labeled AF-1 or Pin1 and samples prepared for ZZ-exchange experiments, containing 400+ µM ^15^N-labeled protein due to the decreased sensitivity of these experiments. Chemical shift assignments previously reported and deposited to the Biological Magnetic Resonance Data Bank (BMRB) for PPARγ AF-1 (43) (BMRB 51507) and Pin1 (62) (BMRB 27579); assignment transfer was confirmed using 3D NMR datasets (e.g., HNCO, HNCA, HNCACO, HNCACB, and HNCOCACB). P76A mutant AF-1 peak assignment was straightforward and did not require 3D NMR experiments of ^13^C,^15^N-labeled mutant protein. Comparison of [^1^H,^15^N]-HSQC spectra of WT and P76A mutant show clear emergence of a new peak corresponding to A76 in a region of the spectra sparsely populated by peaks, where the nearest peaks correspond to residues distant from the mutation site. Peaks nearby A76 experience chemical shift perturbations which scale in intensity with their protein position relative to the mutation site. Titration analysis of ^15^N Pin1 with AF-1 peptides, unphosphorylated AF-1, phosphorylated AF-1 (pAF-1) or phosphorylated pAF-1 P76A mutant was followed using 2D [^1^H,^15^N]-TROSY-HSQC experiments. Titration analysis of ^15^N AF-1 (wild-type/PF76A mutant, phosphorylated and unphosphorylated) with Pin1 (full-length, WW domain, or PPIase domain) was followed using 2D [^1^H,^15^N]-HSQC experiments. Concentrations of unlabeled binding partner in solution of titration studies is as follows: ^15^N Pin1 (200 µM) + peptides (pS112, S112, pT75, T75, W39); 0.25X, 0.5X, 0.75X, 1X, 2X, 3X, 4X, 5X, ^15^N AF-1 (200 µM) + Pin1; 0.2X, 0.4X, 0.6X, 0.8X, 1X, 1.5X, 2X, 2.5X, 3X, ^15^N pAF-1 (200 µM) + Pin1; 0.2X, 0.4X, 0.6X, 0.8X, 1X, 1.5X, 2X, 2.5X, 3X, ^15^N Pin1 (200 µM) + AF-1; 0.25X, 0.5X, 0.75X, 1X, 1.5X, 2X, ^15^N Pin1 (200 µM) + pAF-1; 0.2X, 0.4X, 0.6X, 0.8X, 1X, 1.5X, 2X. Chemical shift perturbation analysis performed using the minimal chemical shift method (50) using the following equation: (^1^H_ppm_^titration^ – ^1^H_ppm_^apo^)^2^ + [α(^15^N_ppm_^titration^ – ^15^N_ppm_^apo^)]^2^ where α is the ratio of the ^15^N and ^1^H gyromagnetic ratios (−27130000 and 267520000 rad s^−1^ T^−1^, respectively). 2D [^1^H,^15^N]-ZZ-exchange NMR data were collected with an exchange delay (d7) of 0.05 or 1.5 s. Paramagnetic relaxation enhancement (PRE) NMR experiments were performed using ^15^N-labeled Pin1 (200 µM) and 0.5 molar equivalent of MTSSL-labeled PPARγ AF-1 mutant protein (D33C,T75A) containing a D33C mutation to attach MTSSL, previously used for PRE NMR studies (43), and T75A mutation to exclusively phosphorylate S112. Proteins were labeled via overnight incubation with MTSSL at 10 molar equivalents. MTSSL-labeled protein was dialyzed overnight to remove excess spin label. PRE NMR 2D [^1^H,^15^N]-TROSY-HSQC experiments were collected in the absence or presence of 5X molar excess of sodium ascorbate to reduce the MTSSL nitroxide spin label; I_PRE_ values were calculated from the ratio of peak intensities (I_para_/I_dia_) after normalizing for dilution by the addition of sodium ascorbate. All NMR data were collected using TopSpin software (Bruker Biospin) and processed/analyzed using NMRFx software (One Moon Scientific) (63). Titration data were fit using NMR TITAN (51, 52).

### NMR TITAN binding affinity calculations

NMR TITAN lineshape analysis software (v1.6) was used for the calculation of binding affinities (***K_D_***) and off-rates (***k_off_***) using titration datasets of ^15^N-labeled Pin1 + pAF-1 and ^15^N-labeled pAF-1 + Pin1. The two-state binding model (*P + L* ⇌ *PL*) was used for the purposes of independently fitting *K_D_* and *k_off_* for the WW domain of Pin1 using a selection of peaks corresponding to amino acids specific to that domain as indicated in **Table 1**. The same two state model was used for the reciprocal calculation with ^15^N-labeled AF-1 using peaks corresponding to amino acids at the pS112 binding site. Datasets of ^15^N-labeled pAF-1 (200 µM) and ^15^N-labeled Pin1 (200 µM) with titrations of unlabeled phosphorylated AF-1 or Pin1, respectively, at 0X, 0.2X, 0.4X, 0.6X, 0.8X, and 1X, were used for these calculations (partial data shown in **Figures 2 and 3**, respectively; and fitted data in ***SI Appendix*, Figs. S5 and S6** and **Datasets S1 and S2**). Upon selection of peaks to be used for fitting, the free (0X) and bound state (1X) chemical shifts and linewidths of select residues were fit and chemical shifts then consequently fixed during the fitting of the remainder of parameters along with the *K_D_* and *k_off_*. Plots of experimental and simulated data can be found in the supporting information, along with the results of bootstrap error analyses performed using 200 replicas.

### Cell culture, luciferase reporter assays, and RT-qPCR

HEK293T cells (ATCC, 12022001) were cultured in Dulbecco’s minimal essential medium (DMEM, Gibco 11960-044) supplemented with 10% fetal bovine serum (FBS, Gibco 26140-079) and Penicillin-Streptomycin-Glutamine (Gibco 10378-016) at 37°C in 5% CO2 atmosphere. 15,000 cells/well were plated in a 96 well plate (Greiner 655083) in 100 µL of media, left overnight, and transfected the following day with 50 ng PPRE, 25 ng PIN1, 25 ng N-terminal HA-tagged full length PPARγ2 (Sino Biological HG12019-NY), and 25 ng pCDNA3.1 empty vector in accordance with experimental conditions using Lipofectamine 3000 (Invitrogen L3000008) and OptiMEM (Gibco 31985-070). The PPRE plasmid is a luciferase reporter which contains three copies of the PPAR-binding DNA response element sequence (3xPPRE-luciferase). Immediately prior to transfection, media in wells was replaced with media supplemented with KPT-6566 or equivalent volume of DMSO in accordance with experimental conditions. After 48h of incubation 25 µL of Bright-Glo reagent (Promega E2620) was added to each well and the luminescence measured using a BioTek Synergy Neo plate reader. For gene expression measurements by RT-qPCR, RNA was extracted from HEK293T cells using Zymo Research Quick-RNA MiniPrep (R1055) according to manufacturer’s protocol. 2 µg of extracted RNA was then reverse transcribed using BioRad iScript cDNA synthesis kit (1708890) according to manufacturer’s protocol. cDNA was diluted by one half for use in qPCR reactions. 10 µL qPCR reactions contained 0.1 µL of 100 µM reverse and forward primers, 4.8 µL of diluted cDNA, and 5 µL of 2X PowerTrack SYBR Green Master Mix (applied biosystems, A46109) and measured in a comparative C_T_ experiment using Applied Biosystems QuantStudio5. PIN1 primer sequence: F-TCAGGCCGAGTGTACTACTTC, R-TCTTCTGGATGTAGCCGTTGA; GAPDH primer sequence: F-TGCACCACCAACTGCTTAGC, R-GGCATGGACTGTGGTCATGAG

## ACKNOWLEDGMENTS

This work was supported in part by National Institutes of Health (NIH) grants R01DK124870 (DJK) and F31DK134167 (CCW) from the National Institute of Diabetes and Digestive and Kidney Diseases (NIDDK) as well as grants for NMR instrumentation from NIH (S10OD021550) to Scripps Florida for the purchase of a 600 MHz NMR; NSF-MRI (0922862) for the acquisition of a 900 MHz Ultra-High Field NMR spectrometer; NIH (S10 RR025677) for console upgrades on all biomolecular NMR spectrometers in 2009; NIH (R35GM118089-04S1), NIH supplement for the helium liquefier; NIH S10OD034276 to replace a 800 MHz spectrometer, accompanied by Vanderbilt University matching funds. The contents of this publication are solely the responsibility of the authors and do not necessarily represent the official views of NIDDK or NIH. We thank George Tsaprailis and the staff at the mass spectrometry and proteomics core at the Herbert Wertheim UF Scripps Institute for Biomedical Innovation and Technology for their aid in the collection and analysis of proteomics data presented herein.

## DATA AVAILABILITY

Mass spectrometry results have been deposited in the massIVE database with the accession number MSV000096300.

## AUTHOR CONTRIBUTIONS

C.C.W. performed all experimental studies. J.C. performed *in vitro* phosphorylation of samples for mass spectrometry analysis. P.M.T. expressed and purified proteins. D.K. supervised the research. C.C.W. and D.J.K. conceived and designed the research, analyzed data, and wrote the manuscript.

## REFERENCES

1. D. E. Stewart, A. Sarkar, J. E. Wampler, Occurrence and role of cis peptide bonds in protein structures. J Mol Biol 214, 253–260 (1990).

2. P. A. Schmidpeter, J. R. Koch, F. X. Schmid, Control of protein function by prolyl isomerization. Biochim Biophys Acta 1850, 1973–1982 (2015).

3. U. Reimer et al., Side-chain effects on peptidyl-prolyl cis/trans isomerisation. J Mol Biol 279, 449–460 (1998).

4. M. Schutkowski et al., Role of phosphorylation in determining the backbone dynamics of the serine/threonine-proline motif and Pin1 substrate recognition. Biochemistry 37, 5566–5575 (1998).

5. I. R. Kleckner, M. P. Foster, An introduction to NMR-based approaches for measuring protein dynamics. Biochim Biophys Acta 1814, 942–968 (2011).

6. H. N. Cheng, F. A. Bovey, Cis-trans equilibrium and kinetic studies of acetyl-L-proline and glycyl-L-proline. Biopolymers 16, 1465–1472 (1977).

7. K. P. Lu, G. Finn, T. H. Lee, L. K. Nicholson, Prolyl cis-trans isomerization as a molecular timer. Nat Chem Biol 3, 619–629 (2007).

8. K. P. Lu, X. Z. Zhou, The prolyl isomerase PIN1: a pivotal new twist in phosphorylation signalling and disease. Nat Rev Mol Cell Biol 8, 904–916 (2007).

9. A. H. Andreotti, Native state proline isomerization: an intrinsic molecular switch. Biochemistry 42, 9515–9524 (2003).

10. F. Zosel, D. Mercadante, D. Nettels, B. Schuler, A proline switch explains kinetic heterogeneity in a coupled folding and binding reaction. Nat Commun 9, 3332 (2018).

11. G. M. Wulf, Y. C. Liou, A. Ryo, S. W. Lee, K. P. Lu, Role of Pin1 in the regulation of p53 stability and p21 transactivation, and cell cycle checkpoints in response to DNA damage. J Biol Chem 277, 47976–47979 (2002).

12. H. Zheng et al., The prolyl isomerase Pin1 is a regulator of p53 in genotoxic response. Nature 419, 849–853 (2002).

13. X. Sun, H. J. Dyson, P. E. Wright, A phosphorylation-dependent switch in the disordered p53 transactivation domain regulates DNA binding. Proc Natl Acad Sci U S A 118 (2021).

14. B. A. Hilton et al., ATR Plays a Direct Antiapoptotic Role at Mitochondria, which Is Regulated by Prolyl Isomerase Pin1. Mol Cell 60, 35–46 (2015).

15. S. C. Lummis et al., Cis-trans isomerization at a proline opens the pore of a neurotransmitter-gated ion channel. Nature 438, 248–252 (2005).

16. S. D. Hanes, Prolyl isomerases in gene transcription. Biochim Biophys Acta 1850, 2017–2034 (2015).

17. T. Hunter, Prolyl Isomerases and Nuclear Function. Cell 92, 141–143 (1998).

18. L. Pastorino et al., The prolyl isomerase Pin1 regulates amyloid precursor protein processing and amyloid-beta production. Nature 440, 528–534 (2006).

19. D. Gurung et al., Proline Isomerization: From the Chemistry and Biology to Therapeutic Opportunities. Biology (Basel) 12 (2023).

20. C. L. Gustafson et al., A Slow Conformational Switch in the BMAL1 Transactivation Domain Modulates Circadian Rhythms. Mol Cell 66, 447–457 e447 (2017).

21. P. Sarkar, C. Reichman, T. Saleh, R. B. Birge, C. G. Kalodimos, Proline cis-trans isomerization controls autoinhibition of a signaling protein. Mol Cell 25, 413–426 (2007).

22. J. Fanghanel, G. Fischer, Insights into the catalytic mechanism of peptidyl prolyl cis/trans isomerases. Front Biosci 9, 3453–3478 (2004).

23. P. E. Shaw, Peptidyl-prolyl isomerases: a new twist to transcription. EMBO Rep 3, 521–526 (2002).

24. C. J. Tsai, B. Ma, R. Nussinov, Protein-protein interaction networks: how can a hub protein bind so many different partners? Trends Biochem Sci 34, 594–600 (2009).

25. P. J. Lu, X. Z. Zhou, M. Shen, K. P. Lu, Function of WW domains as phosphoserine- or phosphothreonine-binding modules. Science 283, 1325–1328 (1999).

26. M. A. Verdecia, M. E. Bowman, K. P. Lu, T. Hunter, J. P. Noel, Structural basis for phosphoserine-proline recognition by group IV WW domains. Nat Struct Biol 7, 639–643 (2000).

27. R. Ranganathan, K. P. Lu, T. Hunter, J. P. Noel, Structural and functional analysis of the mitotic rotamase Pin1 suggests substrate recognition is phosphorylation dependent. Cell 89, 875–886 (1997).

28. K. P. Lu, S. D. Hanes, T. Hunter, A human peptidyl-prolyl isomerase essential for regulation of mitosis. Nature 380, 544–547 (1996).

29. M. B. Yaffe et al., Sequence-specific and phosphorylation-dependent proline isomerization: a potential mitotic regulatory mechanism. Science 278, 1957–1960 (1997).

30. A. Born et al., Ligand-specific conformational change drives interdomain allostery in Pin1. Nat Commun 13, 4546 (2022).

31. J. Chen, A Specific pSer/Thr-Pro Motif Generates Interdomain Communication Bifurcations of Two Modes of Pin1 in Solution Nuclear Magnetic Resonance. Biochemistry 61, 1167–1180 (2022).

32. X. Wang, B. J. Mahoney, M. Zhang, J. S. Zintsmaster, J. W. Peng, Negative Regulation of Peptidyl-Prolyl Isomerase Activity by Interdomain Contact in Human Pin1. Structure 23, 2224–2233 (2015).

33. J. W. Peng, Investigating Dynamic Interdomain Allostery in Pin1. Biophys Rev 7, 239–249 (2015).

34. T. Mori, S. Saito, Molecular Insights into the Intrinsic Dynamics and Their Roles During Catalysis in Pin1 Peptidyl-prolyl Isomerase. J Phys Chem B 126, 5185–5193 (2022).

35. M. Zhang, T. E. Frederick, J. VanPelt, D. A. Case, J. W. Peng, Coupled intra- and interdomain dynamics support domain cross-talk in Pin1. J Biol Chem 295, 16585–16603 (2020).

36. R. La Montagna, I. Caligiuri, A. Giordano, F. Rizzolio, Pin1 and nuclear receptors: a new language? J Cell Physiol 228, 1799–1801 (2013).

37. T. P. Burris et al., Nuclear receptors and their selective pharmacologic modulators. Pharmacol Rev 65, 710–778 (2013).

38. M. I. Lefterova, A. K. Haakonsson, M. A. Lazar, S. Mandrup, PPARgamma and the global map of adipogenesis and beyond. Trends Endocrinol Metab 25, 293–302 (2014).

39. Y. Fujimoto et al., Proline cis/trans-isomerase Pin1 regulates peroxisome proliferator-activated receptor gamma activity through the direct binding to the activation function-1 domain. J Biol Chem 285, 3126–3132 (2010).

40. Y. Han, S. H. Lee, M. Bahn, C. Y. Yeo, K. Y. Lee, Pin1 enhances adipocyte differentiation by positively regulating the transcriptional activity of PPARgamma. Mol Cell Endocrinol 436, 150–158 (2016).

41. P. Rajagopal, E. B. Waygood, J. Reizer, M. H. Saier, Jr., R. E. Klevit, Demonstration of protein-protein interaction specificity by NMR chemical shift mapping. Protein Sci 6, 2624–2627 (1997).

42. D. A. Bosco, E. Z. Eisenmesser, S. Pochapsky, W. I. Sundquist, D. Kern, Catalysis of cis/trans isomerization in native HIV-1 capsid by human cyclophilin A. Proc Natl Acad Sci U S A 99, 5247–5252 (2002).

43. S. A. Mosure et al., Structural basis of interdomain communication in PPARγ. bioRxiv 10.1101/2022.07.13.499031, 2022.2007.2013.499031 (2022).

44. M. Adams, M. J. Reginato, D. Shao, M. A. Lazar, V. K. Chatterjee, Transcriptional activation by peroxisome proliferator-activated receptor gamma is inhibited by phosphorylation at a consensus mitogen-activated protein kinase site. J Biol Chem 272, 5128–5132 (1997).

45. J. H. Choi et al., Anti-diabetic drugs inhibit obesity-linked phosphorylation of PPARgamma by Cdk5. Nature 466, 451–456 (2010).

46. A. S. Banks et al., An ERK/Cdk5 axis controls the diabetogenic actions of PPARgamma. Nature 517, 391–395 (2015).

47. F. A. Gonzalez, D. L. Raden, R. J. Davis, Identification of substrate recognition determinants for human ERK1 and ERK2 protein kinases. J Biol Chem 266, 22159–22163 (1991).

48. M. Cargnello, P. P. Roux, Activation and function of the MAPKs and their substrates, the MAPK-activated protein kinases. Microbiol Mol Biol Rev 75, 50–83 (2011).

49. R. Hendus-Altenburger et al., Random coil chemical shifts for serine, threonine and tyrosine phosphorylation over a broad pH range. J Biomol NMR 73, 713–725 (2019).

50. M. P. Williamson, Using chemical shift perturbation to characterise ligand binding. Prog Nucl Magn Reson Spectrosc 73, 1–16 (2013).

51. C. A. Waudby, J. Christodoulou, NMR Lineshape Analysis of Intrinsically Disordered Protein Interactions. Methods Mol Biol 2141, 477–504 (2020).

52. C. A. Waudby, A. Ramos, L. D. Cabrita, J. Christodoulou, Two-Dimensional NMR Lineshape Analysis. Sci Rep 6, 24826 (2016).

53. E. Campaner et al., A covalent PIN1 inhibitor selectively targets cancer cells by a dual mechanism of action. Nat Commun 8, 15772 (2017).

54. G. Franciosa et al., Prolyl-isomerase Pin1 controls Notch3 protein expression and regulates T-ALL progression. Oncogene 35, 4741–4751 (2016).

55. Y. M. Lee, Y. C. Liou, Gears-In-Motion: The Interplay of WW and PPIase Domains in Pin1. Front Oncol 8, 469 (2018).

56. B. T. Innes, M. L. Bailey, C. J. Brandl, B. H. Shilton, D. W. Litchfield, Non-catalytic participation of the Pin1 peptidyl-prolyl isomerase domain in target binding. Front Physiol 4, 18 (2013).

57. M. J. Rogals, A. I. Greenwood, J. Kwon, K. P. Lu, L. K. Nicholson, Neighboring phosphoSer-Pro motifs in the undefined domain of IRAK1 impart bivalent advantage for Pin1 binding. FEBS J 283, 4528–4548 (2016).

58. S. Hadzi, R. Loris, J. Lah, The sequence-ensemble relationship in fuzzy protein complexes. Proc Natl Acad Sci U S A 118 (2021).

59. E. Komives, R. Sanchez-Rodriguez, H. Taghavi, M. Fuxreiter, Fuzzy protein-DNA interactions and beyond: A common theme in transcription? Curr Opin Struct Biol 89, 102941 (2024).

60. M. P. Williamson, Protein Binding: A Fuzzy Concept. Life (Basel) 13 (2023).

61. J. A. Purslow, B. Khatiwada, M. J. Bayro, V. Venditti, NMR Methods for Structural Characterization of Protein-Protein Complexes. Front Mol Biosci 7, 9 (2020).

62. A. Born et al., Backbone and side-chain chemical shift assignments of full-length, apo, human Pin1, a phosphoprotein regulator with interdomain allostery. Biomol NMR Assign 13, 85–89 (2019).

63. M. Norris, B. Fetler, J. Marchant, B. A. Johnson, NMRFx Processor: a cross-platform NMR data processing program. J Biomol NMR 65, 205–216 (2016).

