## Supplementary Figures S1-S6 for "A tethering mechanism underlies Pin1-catalyzed proline *cis-trans* isomerization at a noncanonical site"

### Supporting Information for
A tethering mechanism underlies Pin1-catalyzed proline *cis-trans* isomerization at a noncanonical site

Christopher C. Williams [1,2], Jonathan Chuck [1,2], Paola Munoz-Tello [3], and Douglas J. Kojetin [2,3,4,5,6,*]

[1] Skaggs Graduate School of Chemical and Biological Sciences at Scripps Research, Jupiter, United States
[2] Department of Integrative Structural and Computational Biology, Scripps Research and The Herbert Wertheim UF Scripps Institute for Biomedical Innovation & Technology, Jupiter, Florida, United States
[3] Department of Biochemistry, Vanderbilt University, Nashville, Tennessee, United States
[4] Center for Structural Biology, Vanderbilt University, Nashville, Tennessee, United States
[5] Vanderbilt Institute of Chemical Biology, Vanderbilt University, Nashville, Tennessee, United States
[6] Center for Applied AI in Protein Dynamics, Vanderbilt University, Nashville, Tennessee, United States
* Corresponding author: Douglas J. Kojetin


**This PDF file includes:**

Figures S1 to S6
Legends to Datasets S1 and S2

**Other supporting materials for this manuscript include the following:**

Datasets S1 and S2

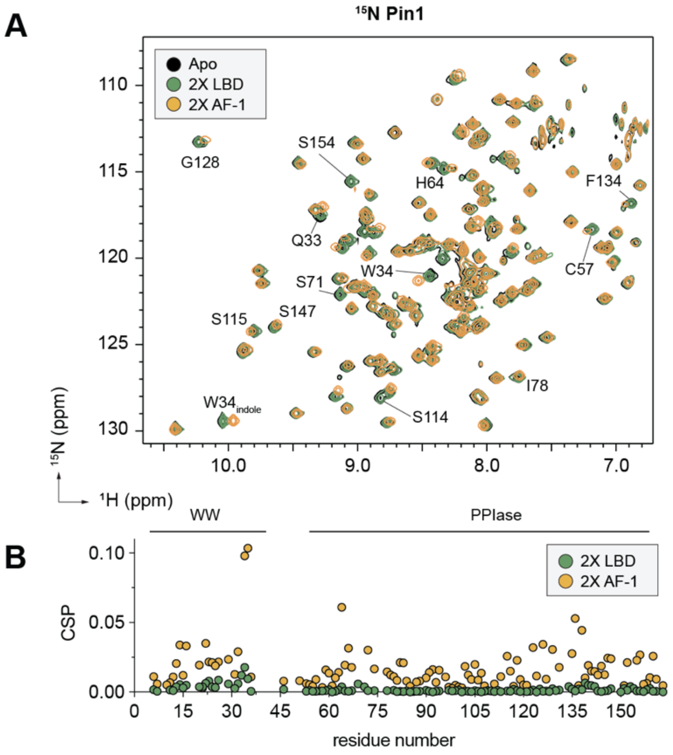


**Fig. S1. 2D [$^{1}$H,$^{15}$N]-TROSY HSQC NMR of $^{15}$N Pin1 + PPARγ2 AF-1 and LBD.** (**A**) $^{15}$N Pin1 (apo, black) + 2X PPARγ2 LBD (green) or 2X AF-1 (mustard) demonstrate that in an unphosphorylated state, the LBD makes no substantial interactions with Pin1, while the AF-1 region makes weak interactions with key residues in both the WW and PPIase domains. (**B**) Quantification of CSPs elicited by 2X LBD and AF-1.

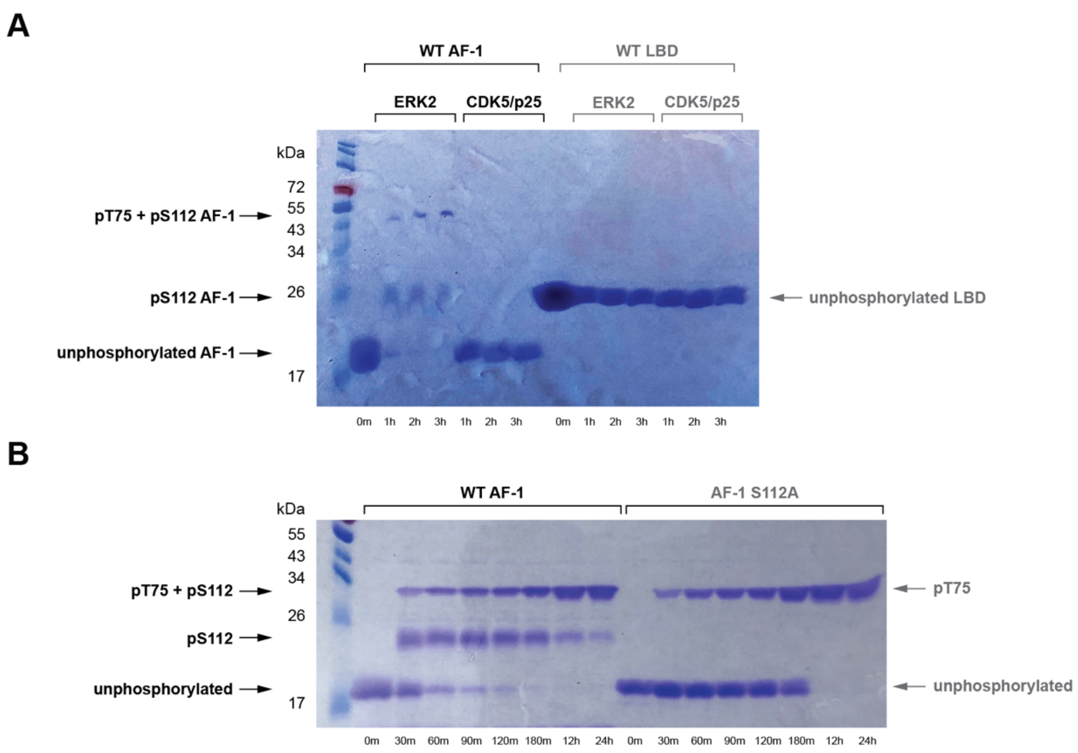


**Fig. S2. Phosphorylation reactions of PPARγ LBD, AF-1 WT, and AF-1 S112A mutant protein followed by Phos-tag SDS-PAGE.** (**A**) ERK2 and CDK/p25 phosphorylation reaction time course with AF-1 and LBD. (**B**) ERK2 phosphorylation reaction time course with AF-1 WT and S112A mutant, which were used for mass spectrometry identification of phosphorylation sites (pS112 and pT75).

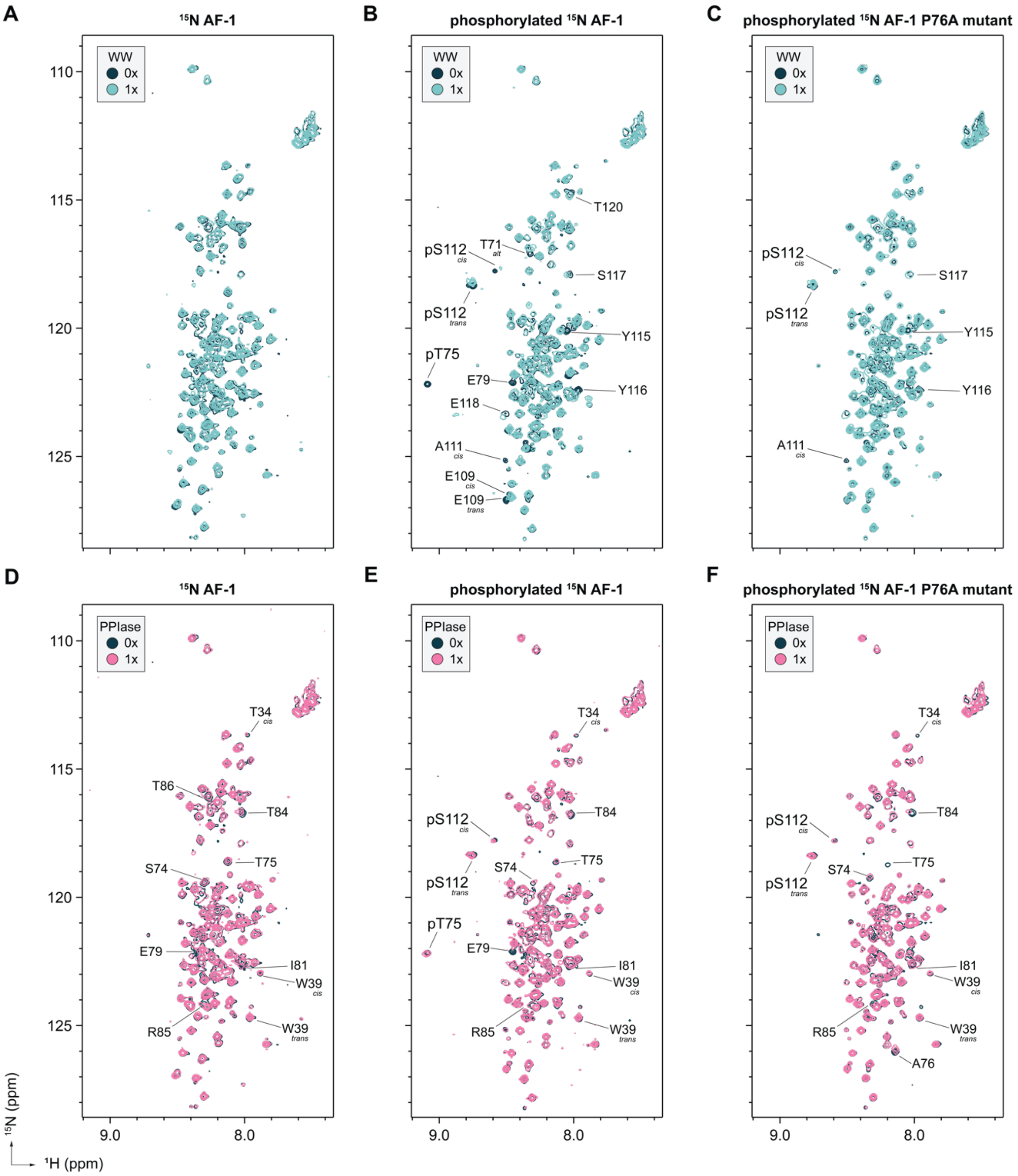


**Fig. S3. Pin1 WW and PPIase domains synergistically enhance binding to phosphorylated AF-1.** Overlays of 2D [$^{1}$H,$^{15}$N]-HSQC NMR spectra of (**A,D**) $^{15}$N-labeled AF-1, (**B,E**) $^{15}$N-labeled pAF-1, and (**C,F**) $^{15}$N-labeled pAF-1 P76A mutant without or with 1 molar equivalent of Pin1 WW domain (**A-C**) or Pin1 PPIase domain (**D-F**).

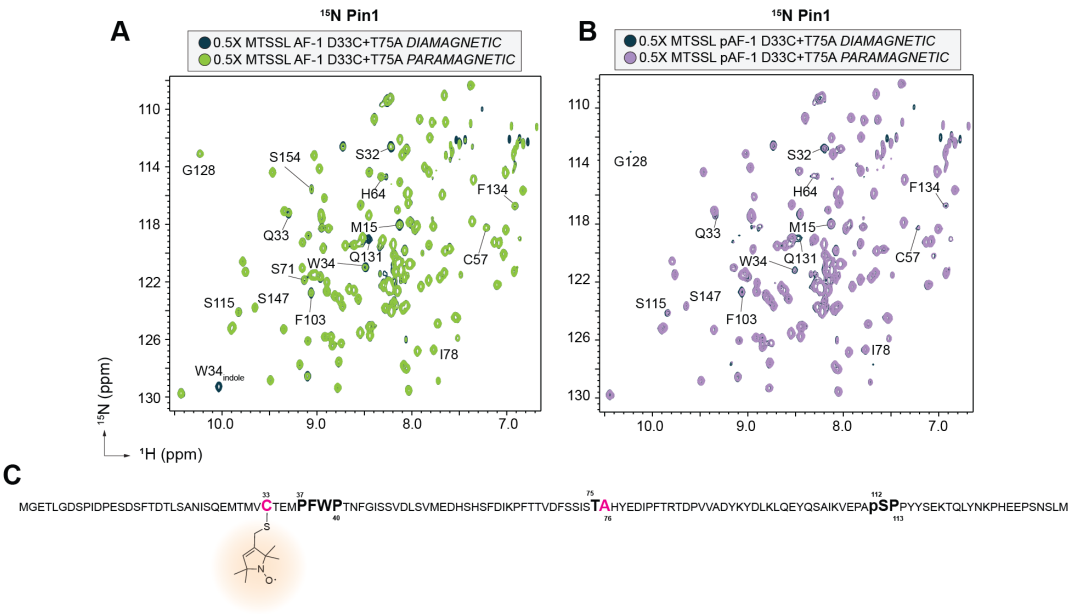


**Fig. S4. Paramagnetic relaxation enhancement (PRE) NMR of $^{15}$N Pin1 with mutant AF-1 constructs.** (**A**) 2D [1H,15N]-TROSY HSQC of $^{15}$N Pin1 + 0.5X MTSSL labeled AF-1 D33C + T75A diamagnetic (navy) and paramagnetic (green) states. (**B**) $^{15}$N Pin1 + 0.5X MTSSL labeled pAF-1 D33C + T75A diamagnetic (navy) and paramagnetic (purple) states. (**C**) Sequence of AF-1 showing sites of mutation (magenta; D33C, T75A), phosphorylation (S112), and MTSSL labeling (D33C).

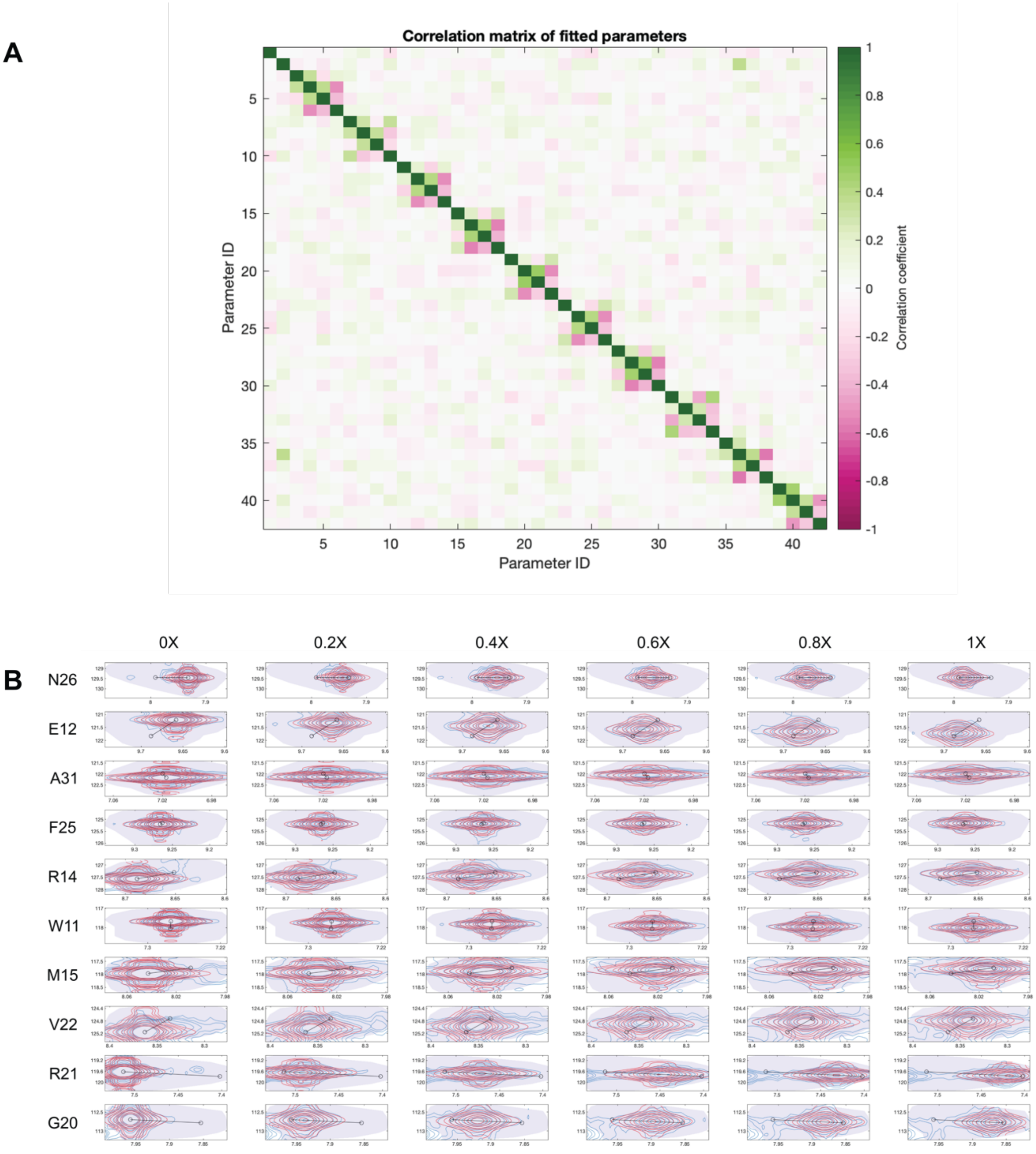
A
Correlation matrix of fitted parameters
Parameter ID
Parameter ID
Correlation coefficient
B
0X
0.2X
0.4X
0.6X
0.8X
1X
N26
E12
A31
F25
R14
W11
M15
V22
R21
G20

**C**

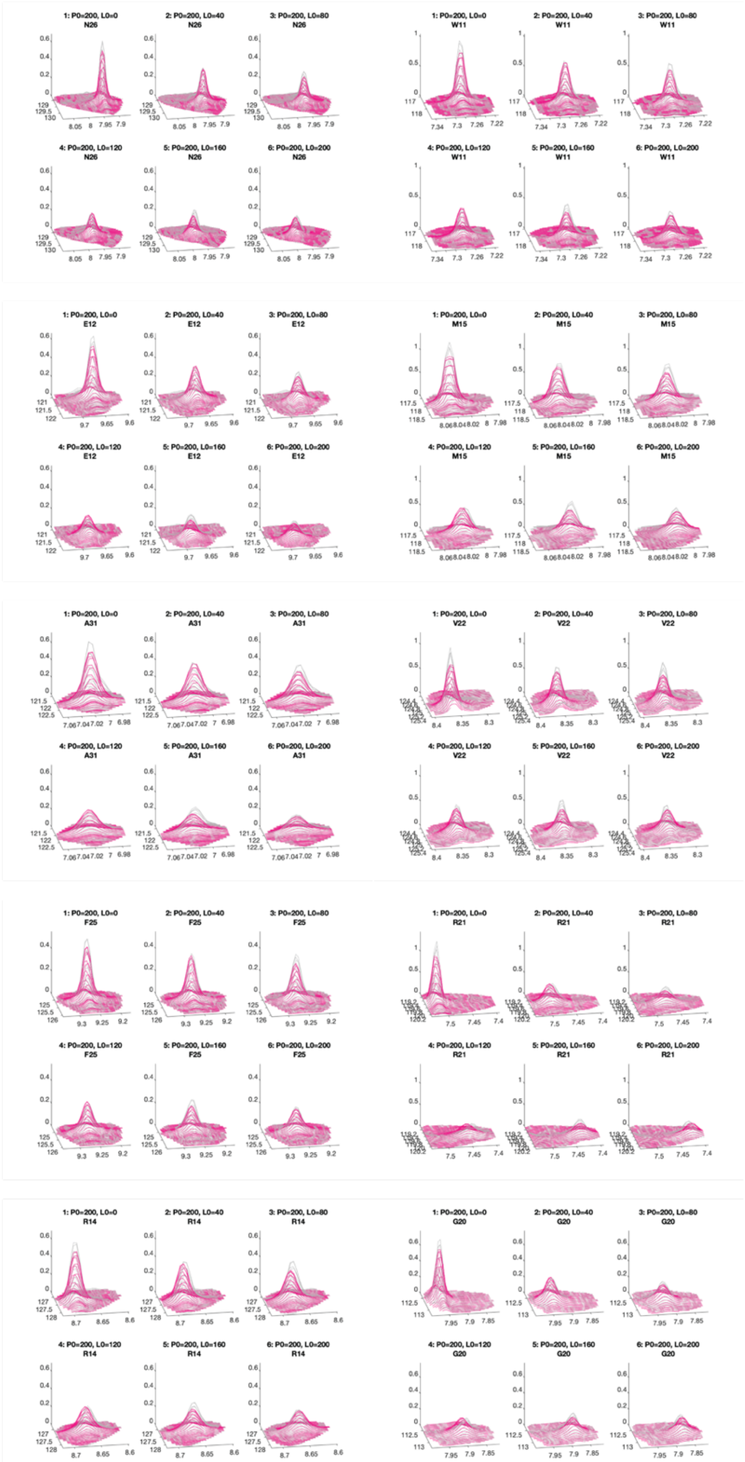

**Fig S5. NMR TITAN $K_D$ fitting results of $^{15}$N-labeled Pin1 titration 2D [$^{1}$H,$^{15}$N]-TROSY HSQC NMR spectra.** (**A**) Covariance plot results of bootstrap analysis performed with 200 replicas. Parameter IDs are identified in DatasetS2.xlsx (**B**) Plotgrid results of simulated (red) and experimental (blue) peaks at 0X, 0.2X, 0.4X, 0.6X, 0.8X, and 1X titration points of peaks corresponding to W11, E12, R14, M15, G20, R21, V22, F25, N26, and A31. (**C**) 3D overlays of simulated (pink) and experimental (gray) peaks at 0X, 0.2X, 0.4X, 0.6X, 0.8X, and 1X titration points of peaks corresponding to W11, E12, R14, M15, G20, R21, V22, F25, N26, and A31.

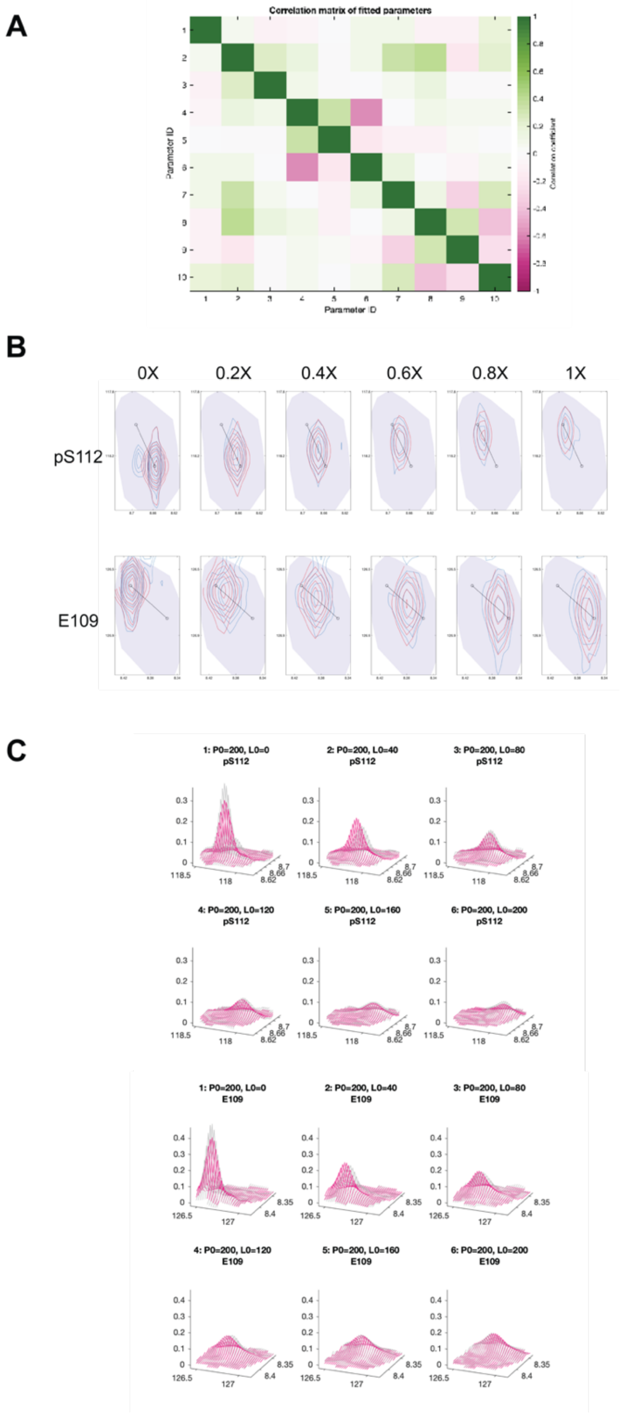


**Fig S6. NMR TITAN $K_D$ fitting results of $^{15}$N-labeled pAF-1 titration 2D [$^{1}$H,$^{15}$N]-HSQC NMR spectra.** (**A**) Covariance plot results of bootstrap analysis performed with 200 replicas. Parameter IDs are identified in DatasetS1.xlsx (**B**) Plotgrid results of simulated (red) and experimental (blue) peaks at 0X, 0.2X, 0.4X, 0.6X, 0.8X, and 1X titration points of peaks corresponding to pS112 and E109. (**C**) 3D overlays of simulated (pink) and experimental (gray) peaks at 0X, 0.2X, 0.4X, 0.6X, 0.8X, and 1X titration points of peaks corresponding to pS112 and E109.

**Dataset S1. Tabulated results of bootstrap analysis of TITAN fitting of $^{15}$N-labeled Pin1 titration data.** Parameter IDs appearing in covariance plots of **Fig. S5** included in this file.

**Dataset S2. Tabulated results of bootstrap analysis of TITAN fitting of $^{15}$N-labeled pAF-1 titration data.** Parameter IDs appearing in covariance plots of **Fig. S6** included in this file.
